# Emerging Drug Combinations for Targeting Tongue Neoplasms Associated Proteins/Genes: Employing Graph Neural Networks within the RAIN Protocol

**DOI:** 10.1101/2024.06.11.598402

**Authors:** Mohsen Askari, Ali A. Kiaei, Mahnaz Boush, Fatemeh Aghaei

## Abstract

**Background:** Tongue Neoplasms is a common form of malignancy, with squamous cell carcinoma of the tongue being the most frequently diagnosed type due to regular mechanical stimulation. Its prevalence remains on the rise among neoplastic cancer cases. Finding effective combinations of drugs to target the genetic and protein elements contributing to the development of Managing Tongue Neoplasms poses a difficulty owing to the intricate and varied nature of the ailment.

**Method:** In this research, we introduce a novel approach using Deep Modularity Networks (DMoN) to identify potential synergistic drug combinations for the condition, following the RAIN protocol. This procedure comprises three primary phases: First, employing Graph Neural Network (GNN) to propose drug combinations for treating the ailment by extracting embedding vectors of drugs and proteins from an extensive knowledge graph containing various biomedical data types, such as drug-protein interactions, gene expression, and drug-target interactions. Second, utilizing natural language processing to gather pertinent articles from clinical trials involving the previously recommended drugs. Finally, conducting network meta-analysis to evaluate the comparative efficacy of these drug combinations.

**Result:** We utilized our approach on a dataset containing drugs and genes as nodes, connected by edges indicating their associated p-values. Our DMoN model identified Cisplatin, Bleomycin, and Fluorouracil as the optimal drug combination for targeting the human genes/proteins associated with this cancer. Subsequent scrutiny of clinical trials and literature confirmed the validity of our findings. Additionally, network meta-analysis substantiated the efficacy of these medications concerning the pertinent genes.

**Conclusion:** Through the utilization of DMoN as part of the RAIN protocol, our method introduces a fresh and effective way to suggest notable drug combinations for addressing proteins/genes linked to Tongue Neoplasms. This approach holds promise in assisting healthcare practitioners and researchers in pinpointing the best treatments for patients, as well as uncovering the fundamental mechanisms of the disease.

**Highlights:**

- A new method using Deep Modularity Networks and the RAIN protocol can find the best drug combinations for treating Tongue Neoplasms, a common and deadly form of cancer.
- The method uses a Graph Neural Network to suggest drug pairings from a large knowledge graph of biomedical data, then searches for clinical trials and performs network meta-analysis to compare their effectiveness.
- The method discovered that Cisplatin, Bleomycin, and Fluorouracil are suitable drugs for targeting the genes/proteins involved in this cancer, and confirmed this finding with literature review and statistical analysis.
- The method offers a novel and powerful way to assist doctors and researchers in finding the optimal treatments for patients with Tongue Neoplasms, and to understand the underlying causes of the disease.

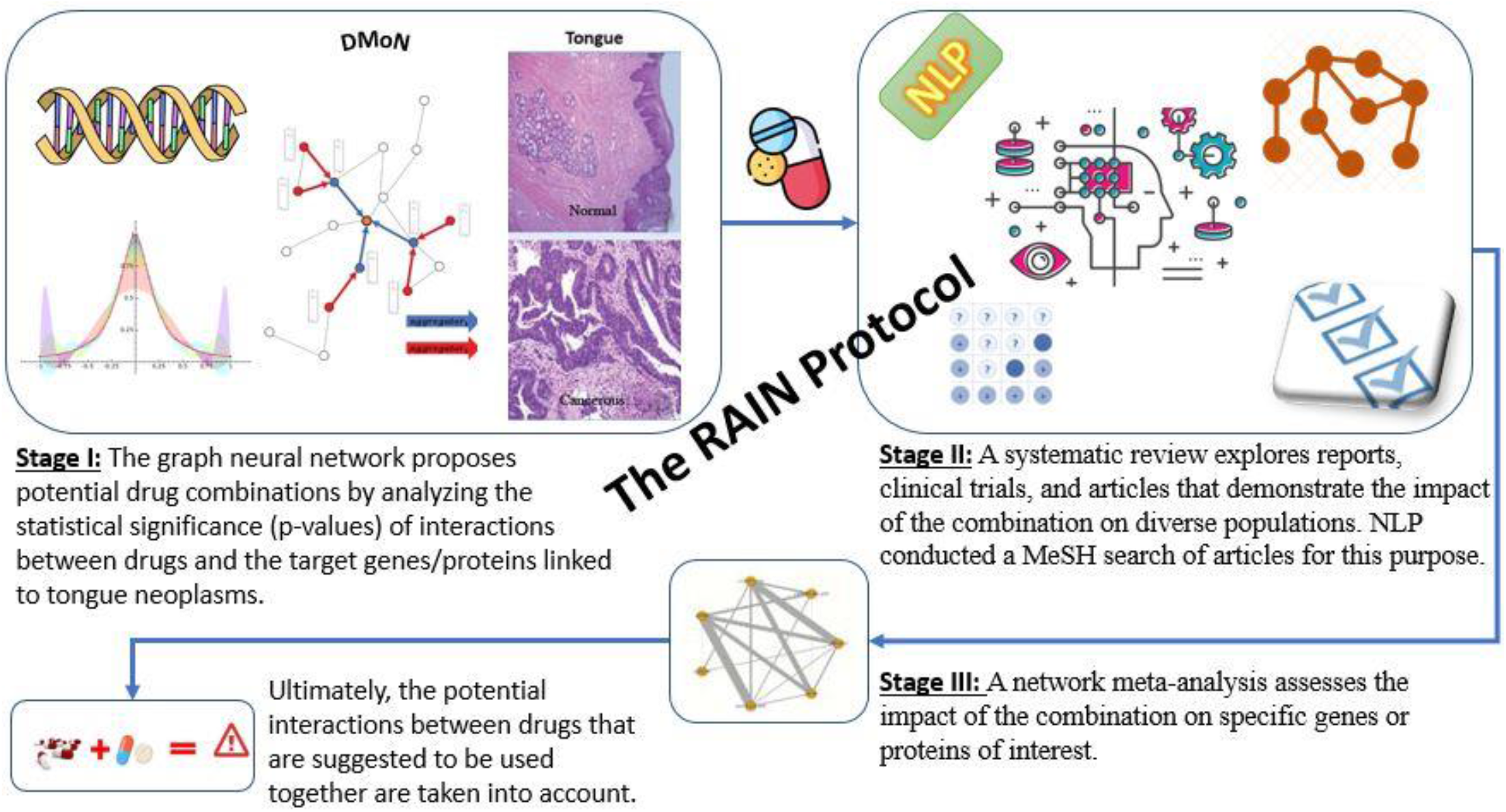

## 1. INTRODUCTION

Tongue neoplasms pertain to the unusual proliferation of cells on the tongue, which may manifest as either benign or malignant. Squamous cell carcinoma stands out as the prevailing form of malignant tongue neoplasm. The outlook for tongue neoplasms is contingent upon factors like cancer stage, tumor size, and the patient’s general well-being. Treatment alternatives encompass surgical intervention, radiation therapy, chemotherapy, or a blend of these approaches. ^1–3^

In the latest release of the World Health Organization’s Classification of Tumours of the Head and Neck, which followed the 2017 fourth edition, significant modifications have been implemented. Specifically, the fifth chapter, titled “Tumours of the oral cavity and mobile tongue,” introduces a new category called Non-neoplastic Lesions, encompassing two additional entries: necrotizing sialo metaplasia and melanoacanthoma. Furthermore, the 2015 WHO section addressing “Oral potentially malignant disorders and Oral epithelial dysplasia” has undergone restructuring, with separate discussions now dedicated to submucous fibrosis and HPV-associated dysplasia. Sections exclusively devoted to carcinoma cuniculatum and verrucous carcinoma have been established, highlighting the oral cavity as the primary site in the head and neck for these conditions. Each of these entities demonstrates distinct clinical and histologic characteristics compared to traditional squamous cell carcinoma. ^3,4^

As per a 2022 publication from the American Association for Cancer Research Reviews, which explores the implications of autonomous drug action in the context of combination therapy within precision oncology, the insights presented in the article are valuable for comprehending the advantages of employing drug combinations in the treatment of tongue neoplasms. [5]

The article examines proof indicating that medication behavior that operates autonomously, initially outlined in 1961, elucidates the effectiveness of numerous composition treatment that have altered medical practices. This concept offers diverse patient populations, characterized by varying drug sensitivities, multiple opportunities to derive benefits from at least one drug. A thorough understanding of varied responses could reveal predictive indicators or biomarkers related to drug effects, allowing for more accurate utilization of current medications and leveraging the benefits of combined or synergistic effects..[6]– [7]

### 1.1 Associated human genes/proteins

X. Liu et al., in their investigation published in BMC Cancer, delved into the role of FAM168A in the progression of chronic myeloid Tongue Neoplasms (CML) through the BCR-ABL1/AKT1/NFκB pathway. The research revealed that within tongue cancer cells, FAM168A, also recognized as tongue cancer resistance-associated protein (TCRP1), has the ability to directly associate with AKT1, influencing the regulation of AKT1/NFκB signaling pathways. Furthermore, the study demonstrated FAM168A’s potential function as a connecting protein, binding both to BCR-ABL1 and AKT1, thereby facilitating downstream signaling pathways in CML^1,8,9^

The genes LncKRT16P6 function as promoters in the advancement of tongue squamous cell carcinoma by acting as sponges for miR-3180 and modulating GATAD2A expression. The study findings demonstrate that LncKRT16P6 boosts the growth, movement, and infiltration of tongue squamous cell carcinoma cells through its interaction with miR-3180, thereby regulating the expression of GATAD2A. Moreover, the research proposes that LncKRT16P6 might represent a promising treatment focus for managing tongue squamous cell carcinoma. ^5–7,10–12^

In the research conducted by D. Hu and D. V. Messadi, which explores the involvement of immune-related long non-coding RNA (lncRNA) signatures in tongue squamous cell carcinoma (TSCC), six immune-related signature lncRNAs, including FNDC1-IT1, were identified with prognostic significance in TSCC. The study determined that the risk score derived from the six-lncRNA model served as a crucial indicator of the survival rate. Additionally, the research illustrated that patient groups classified as high-risk and low-risk exhibited notable distinctions in their immune status. ^13^

A case report by A. Aksionau et al. highlights lingual alveolar soft part sarcoma (ASPS) in a 78-year-old woman. The research identified a specific gene fusion, ASPSCR1 and TFE3, unique to ASPS. The study concludes that ASPS should be considered in the differential diagnoses for slow-growing lingual masses, particularly those with vascular characteristics and nonspecific clinical presentations, irrespective of the patient’s age. ^14,15^

In a different publication, a case report is presented concerning a patient diagnosed with oral squamous cell carcinoma carrying a CDKN2A c.301G>T mutation. The authors explore the heightened susceptibility to head and neck squamous cell carcinoma (HNSCC), specifically oral squamous cell carcinoma (OSCC), in individuals with inherited CDKN2A mutations. They propose that testing for CDKN2A mutations is advisable in cases of familial HNSCC and among young patients lacking apparent risk factors. Furthermore, regular surveillance for HNSCC is recommended as standard practice for individuals with a germline CDKN2A mutation.^16^

The article explores the involvement of CSNK1D in controlling the Wnt/β-catenin signaling pathway, a process implicated in the progression of several cancers, including tongue squamous cell carcinoma (TSCC). The authors propose that directing interventions towards the Wnt/β-catenin signaling pathway could offer a potentially effective therapeutic approach for managing TSCC.^17^

Mutations in TP53 play a role in myeloid malignancies. These publications assert that TP53 mutations hinder the cellular ability to respond to genotoxic stress and instigate inherent resistance to standard cytotoxic treatments. Patients with myeloid malignancies harboring TP53 mutations experience unfavorable clinical outcomes, characterized by high-risk clinical attributes like complex karyotype and previous exposure to leukemogenic therapies. Their survival is typically shortened due to a heightened risk of relapse following allogeneic transplantation. ^5–7,18–20^

The CASC18 gene shows heightened expression in tumors linked to tongue squamous cell carcinoma (TSCC), especially in instances where patients experience occult lymph node metastasis (OLNM). The study revealed a new regulatory axis, CASC18/miR-20a-3p/TGFB2 ceRNA, associated with OLNM in TSCC. These findings have the potential to advance our understanding of the molecular mechanisms involved in OLNM in TSCC and could contribute to the development of diagnostic strategies to assist in treatment decision-making^21^.

J. Li and colleagues conducted a study examining the mRNA expression levels of CDC6, CDT1, MCM2, and CDC45 using quantitative real-time PCR on formalin-fixed, paraffin-embedded benign and malignant tongue tissues. Following this, statistical analysis was conducted. The study unveiled a significant elevation in the expression levels of these genes in malignant squamous cell carcinoma (SCC) compared to mild precancerous epithelial dysplasia. Furthermore, a consistent trend of heightened expression levels corresponding to the severity of precancerous lesions, spanning from mild to moderate and severe epithelial dysplasia, was observed. Relationships were established between CDC6 and CDC45 expression and dysplasia grade and lymph node status. CDT1 expression exhibited an increase in severe dysplasia compared to mild and moderate dysplasia. Dysplasia grade, lymph node status, and clinical stage influenced MCM2 expression. Notably, the expression of these genes was found to be unrelated to tumor size or histological grade. The study concludes that these proteins could serve as biomarkers for distinguishing precancerous dysplasia from SCC, aiding in early detection and diagnosis, particularly when considered alongside clinicopathological parameters. ^22^

SPRR2G is a gene encoding a protein expected to participate in keratinization, situated in both the cornified envelope and cytosol. This protein functions as a cross-linked envelope protein in keratinocytes, initially appearing in the cell cytosol and later becoming cross-linked to membrane proteins by transglutaminase. This process culminates in the creation of an insoluble envelope beneath the plasma membrane. According to the Human Protein Atlas, SPRR2G exhibits expression in diverse cancer tissues, including tongue cancer. ^23, 24^

As of a 2022 article addressing MTFR1’s involvement in tongue squamous cell carcinoma (TSCC), the research indicates that MTFR1 exhibits irregular expression in head and neck squamous cell carcinoma (HNSC), and its specific role in TSCC remains unclear. The study aimed to clarify MTFR1’s function in TSCC by assessing the expression levels of long non-coding RNA small nucleolar RNA host gene 1 (SNHG1), microRNA-194-5p, and MTFR1 in TSCC cells using RT-qPCR. Western blot analysis was employed to scrutinize MTFR1 protein levels and epithelial-mesenchymal transition (EMT) markers. The results demonstrated increased MTFR1 expression in TSCC. Knocking down MTFR1 hindered TGFβ1-induced EMT, along with the migration and invasion of TSCC cells in vitro. MiR-194-5p targeted MTFR1, negatively regulating its expression. Additionally, SNHG1 heightened MTFR1 expression by binding to miR-194-5p. Importantly, SNHG1 stimulated EMT, invasion, and migration of TSCC cells by upregulating MTFR1. The SNHG1/miR-194-5p/MTFR1 axis was identified as a promoter of TGFβ1-induced EMT, migration, and invasion in TSCC cells, suggesting potential targets for treating TSCC patients.^25^

Other review articles have been published that offer a comprehensive overview of immune targets in tongue cancer. These articles delve into the prognostic significance of immune targets associated with tongue cancer and their potential as therapeutic focal points. Additionally, the review underscores the influence of major molecules, linked signaling pathways, epigenetic alterations, DNA damage repair systems, cancer stem cells, and microRNAs in the progression of tongue cancer. MMP9, owing to its proteolytic activity, assumes a pivotal role in tumorigenesis by governing processes such as migration, epithelial-to-mesenchymal transition, cancer cell survival, immune response induction, angiogenesis, and the establishment of the tumor microenvironment. It emerges as an appealing target for anticancer therapies. However, the intricate regulatory mechanisms governing MMP9’s expression, synthesis, and activation, which dictate its specific yet occasionally conflicting roles, along with its high homology with other MMP family members, pose significant challenges in developing effective and safe MMP9 inhibitors as anticancer drugs. Notably, there is optimism surrounding blocking antibodies selectively targeting MMP9, currently undergoing clinical trials. ^26–29^

#### Targeted therapy

Targeted therapy represents a pivotal approach aimed at enhancing overall survival while mitigating undesirable side effects in cancer treatment. This method involves employing drugs designed to specifically target proteins or genes crucial for the growth and dissemination of cancer cells. Typically, these drugs consist of small molecules or biologics that disrupt the growth and division of cancer cells. The specific mechanism of action varies based on the particular protein or gene being targeted.^30, 31^

Cetuximab, a biotechnological medication employed in targeted therapy for tongue neoplasms, functions by attaching to the epidermal growth factor receptor (EGFR) located on the surface of cancer cells. This activity significantly hinders the transmissions that aid in fostering the development of cancerous cells.^32^

Cetuximab-IRDye is a biotechnological treatment utilized in targeted therapy, combining Cetuximab, a monoclonal antibody targeting the epidermal growth factor receptor (EGFR), with IRDye 800CW, a fluorescent dye aiding surgeons in visualizing cancerous tissue during surgical proceduresVia its interaction with EGFR, Cetuximab obstructs the receptor’s ability to convey messages that encourage the proliferation and splitting of malignant cells. Simultaneously, IRDye 800CW assists surgeons in identifying cancerous tissue by emitting light upon exposure to near-infrared light.^33,34^

Copper Cu 64-DOTA-AE105, a biotechnological medication employed in targeted therapy, functions by attaching to uPAR (urokinase-type plasminogen activator receptor), a protein overexpressed in various cancers, including tongue neoplasms. Through this binding process, Copper Cu 64-DOTA-AE105 selectively delivers radiation to cancer cells while preserving normal cells. This characteristic renders it an appealing option for cancer treatment, potentially minimizing the side effects commonly associated with traditional chemotherapy and radiation therapy. ^35,36^

PKI166, classified as a specific therapeutic agent considered a minor compound, functions as an inhibitor of EGFR, HER1, and HER2 proteins vital for the growth and division of cancerous cells. Through the suppression of these proteins, PKI166 has the capacity to impede or halt the proliferation of malignant cells.^37,38^

#### Chemotherapy

Cisplatin, employed as a chemotherapy drug for the treatment of tongue neoplasms, is a small molecule that operates by disrupting the DNA of cancer cells, thereby hindering their ability to divide and proliferate.^39,40^

Bleomycin, employed as a chemotherapy agent for the treatment of tongue neoplasms, is a small molecule that operates by causing damage to the DNA of cancer cells, effectively impeding their ability to divide and proliferate.^41^ Pingyangmycin is a chemotherapy drug classified as a small molecule. Its mechanism of action involves inducing DNA damage and inhibiting DNA synthesis within cancer cells. Pingyangmycin effectively restrains the growth and division of cancer cells by obstructing DNA synthesis^42^.

In a recently published article discussing the application of 1,4-Phenylenebis(Methylene) Selenocyanate in treating tongue neoplasms, this small molecule drug has been utilized in chemoprevention. Its mode of action involves diminishing DNA damage and suppressing mutagenesis within cancer cells. By restraining mutagenesis, 1,4-Phenylenebis(Methylene) Selenocyanate holds promise in obstructing the creation of cancerous cells.^43^

Carboplatin, employed as a chemotherapy medication for various cancer types, including tongue neoplasms, is administered intravenously. Its mechanism of action involves disrupting the DNA within cancer cells, thereby impeding their ability to divide and proliferate. ^44,45^

In the publication authored by M. Zhang et al., it is mentioned that Methotrexate serves as a chemotherapy medication for various cancer types, including tongue neoplasms. It can be administered either intravenously or orally and operates by disrupting the DNA within cancer cells, thus hindering their capacity to divide and proliferate.^46^

#### Immunotherapy

The 2021 publication by U. Majeed and colleagues highlights immunotherapy as a distinct cancer treatment method utilizing the body’s immune system to combat cancer. Immunotherapy drugs function by activating the immune system, prompting it to identify and combat cancer cells. These drugs are also applied in the treatment of tongue neoplasms. One such drug used for tongue neoplasms is Pembrolizumab, a small molecule employed in immunotherapy. Pembrolizumab operates by obstructing the interaction between the programmed death-1 (PD-1) receptor on T cells and its ligands, PD-L1 and PD-L2, on cancer cells. This action enables the immune system to recognize and target cancer cells .^32^

#### Radiotherapy

Fluorodeoxyglucose F18 serves as a radioactive tracer employed in positron emission tomography (PET) scans for the identification of cancer cells within the body. Following injection into the body, the tracer accumulates in regions exhibiting elevated metabolic activity, notably in cancer cells. Subsequently, the PET scan captures the radiation emitted by the tracer, generating images of the body that prove valuable in diagnosing and staging cancer.^47,48^

#### Other treatment strategies

As per recently published articles, H3B-6527 is a small molecule inhibitor with the ability to selectively bind to and obstruct FGFR4. This action hinders the activation of FGFR4, suppresses FGFR4-mediated signaling, and results in the inhibition of cell proliferation in tumor cells that overexpress FGFR4.^49^

Temoporfin, categorized as a minor compound, is utilized in photodynamic therapy (PDT), a cancer treatment approach that utilizes a photosensitizing agent, such as Temoporfin, in combination with a precise wavelength of light to eradicate cancerous cells. Upon exposure to light, Temoporfin generates a type of oxygen that has the capability to eradicate neighboring cancer cells.^50^

### 1.2 Objective

Several recent papers suggest the implementation of the RAIN protocol for disease treatment, which involves combining drugs to bring the p-value closer to one between the illness and target proteins/genes. These studies utilize diverse Artificial Intelligence techniques, such as Graph Neural Network or Reinforcement Learning, to suggest appropriate drug combinations. Typically, they conduct a network meta-analysis to assess comparative efficacy. ^51–62^

The RAIN protocol operates as a systematic review and meta-analysis approach, utilizing the STROBE methodology to investigate a specific medical inquiry. As seen in several recently published medical AI papers, the RAIN protocol stands out by employing AI to tackle a particular medical query. ^63–72^

## 2. METHOD

Finding effective drug combinations for complex diseases is a challenging task, as it requires considering many factors such as the molecular mechanisms of the disease, the pharmacological properties of the drugs, and the clinical evidence of their efficacy and safety. Moreover, there are often many possible drug combinations to choose from, and testing them all experimentally is costly and time-consuming.

We utilized DMoN, a form of Graph Neural Network, to suggest combinations of drugs that are probable to exhibit synergistic effects on the illness. DMoN is capable of learning from an extensive knowledge graph comprising biomedical data of diverse types, including interactions between drugs and proteins, gene expressions, and interactions between drugs and targets. The system has the capability to create embedding vectors to depict drugs and proteins by utilizing the knowledge graph. These vectors are subsequently employed to assess the likeness and structure of various drug pairings. Modularity, a metric measuring how well a graph is clustered, serves as an indicator of the potential synergy of drug combinations.

Through the utilization of DMoN, we’ve compiled a roster of drug combinations with elevated modularity scores, indicating their potential to target identical or correlated genes/proteins implicated in the disease. Nevertheless, this alone doesn’t guarantee the effectiveness and safety of these drug pairings in practical application. Hence, we employed natural language processing to explore clinical trials incorporating the suggested drug pairings and to extract pertinent details including outcomes, dosages, and side effects. This helped us filter out the drug combinations that have negative or conflicting results in clinical settings.

In conclusion, network meta-analysis was employed to evaluate the efficacy of the remaining drug combinations and to categorize them based on their effectiveness on the target genes/proteins. Network meta-analysis is a statistical technique that can combine direct and indirect evidence from multiple sources, and provide a comprehensive and consistent evaluation of different interventions. By using network meta-analysis, we have determined which drug combinations are the most suitable for treating the disease, and provide confidence intervals and ranking probabilities for them.

### 2.1 Stage I: Graph Neural Network

Deep Modularity Networks (DMoN) is a type of neural network architecture designed to facilitate modular learning, where the network learns to decompose a complex task into simpler, more manageable subtasks. This approach is inspired by the concept of modularity in cognitive science, where complex systems are broken down into smaller, more easily understandable modules.

The main idea behind DMoN is to promote better generalization, interpretability, and transferability of learned representations by enforcing modularity within the network. This is achieved through various mechanisms such as explicit modular structure in the network architecture, modular loss functions, and modular training strategies.

Here’s a general overview of how DMoN work:

1. Modular Architecture: DMoN typically consist of multiple interconnected modules, each responsible for learning specific aspects of the task. These modules can be organized hierarchically or in parallel, depending on the nature of the problem.
2. Modular Loss Functions: Instead of optimizing a single global loss function for the entire network, DMoN use modular loss functions that encourage each module to focus on its designated subtask. This helps prevent interference between modules and promotes specialization.
3. Modular Training: DMoN are trained using techniques that promote modularity, such as curriculum learning, where the network is gradually exposed to increasingly complex tasks, or multi-task learning, where the network learns to perform multiple related tasks simultaneously.
4. Inter-module Communication: While each module in a DMN is specialized in its respective subtask, they often need to communicate with each other to share relevant information. This can be achieved through mechanisms such as attention mechanisms or message passing between modules.

Formulas for DMoN can vary greatly depending on the specific architecture and training strategy used. However, some common elements may include:

#### Loss Functions

Each module in a DMoN typically has its own loss function, which may be a combination of standard loss functions such as cross-entropy for classification tasks or mean squared error for regression tasks, along with regularization terms to promote modularity.

#### Forward Propagation

The forward propagation in DMoN involves passing input data through the network’s modules, with each module performing its designated computation. The output of each module is then combined to produce the final output of the network.

#### Backpropagation

During training, the gradients of the modular loss functions with respect to the network parameters are computed using backpropagation. These gradients are then used to update the parameters of each module using optimization algorithms such as stochastic gradient descent (SGD) or variants like Adam.

It’s important to note that DMoN are a broad class of neural network architectures, and there isn’t a single set of formulas that applies to all DMoN. The specific formulas and techniques used depend on the particular implementation and the problem being addressed. The suggested model can be observed as a graphical depiction of this procedure in Figure 3.

**Figure 1:**
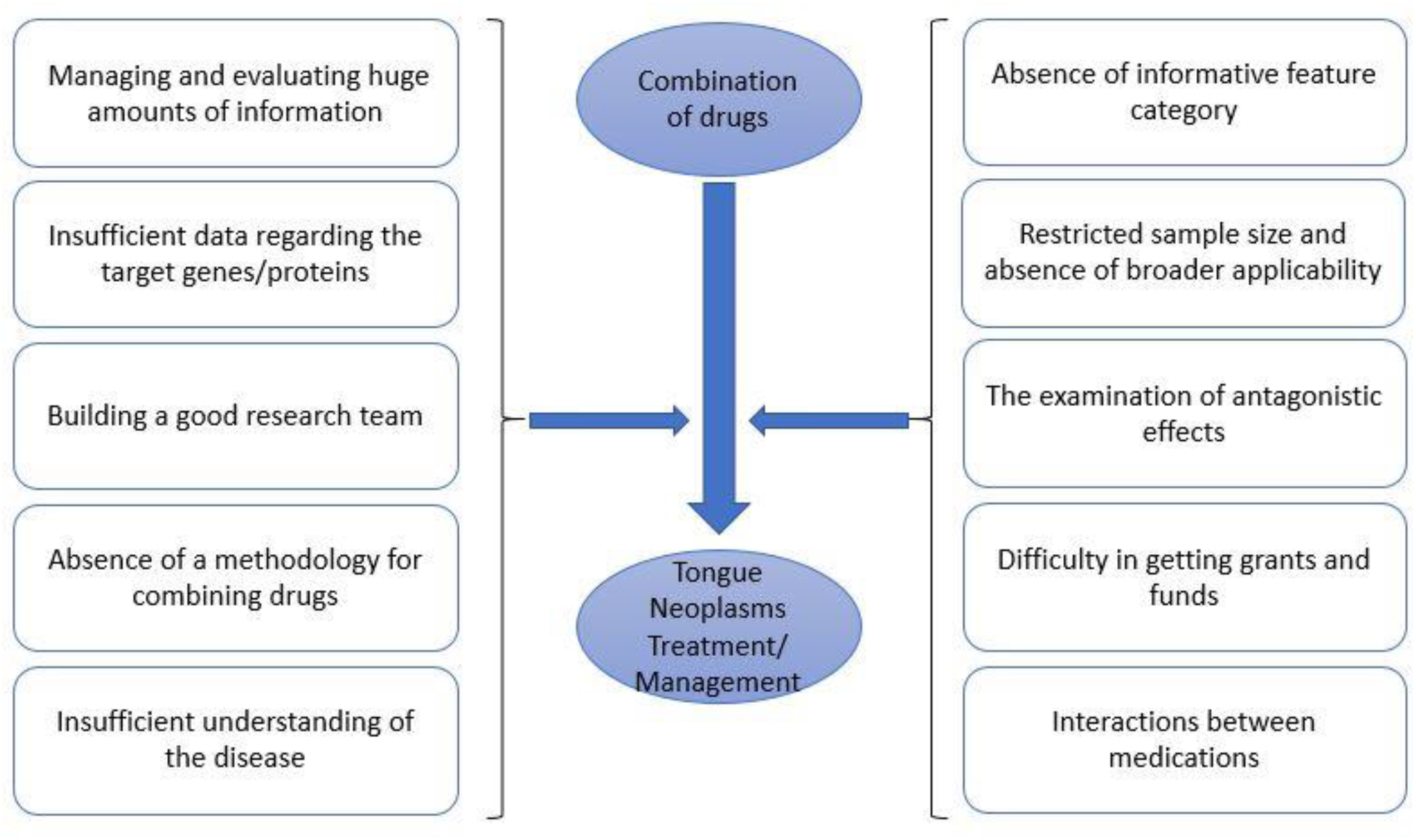
The impacts of suggested drug combinations on the treatment of occurrences of Tongue Neoplasms.

**Figure 2:**
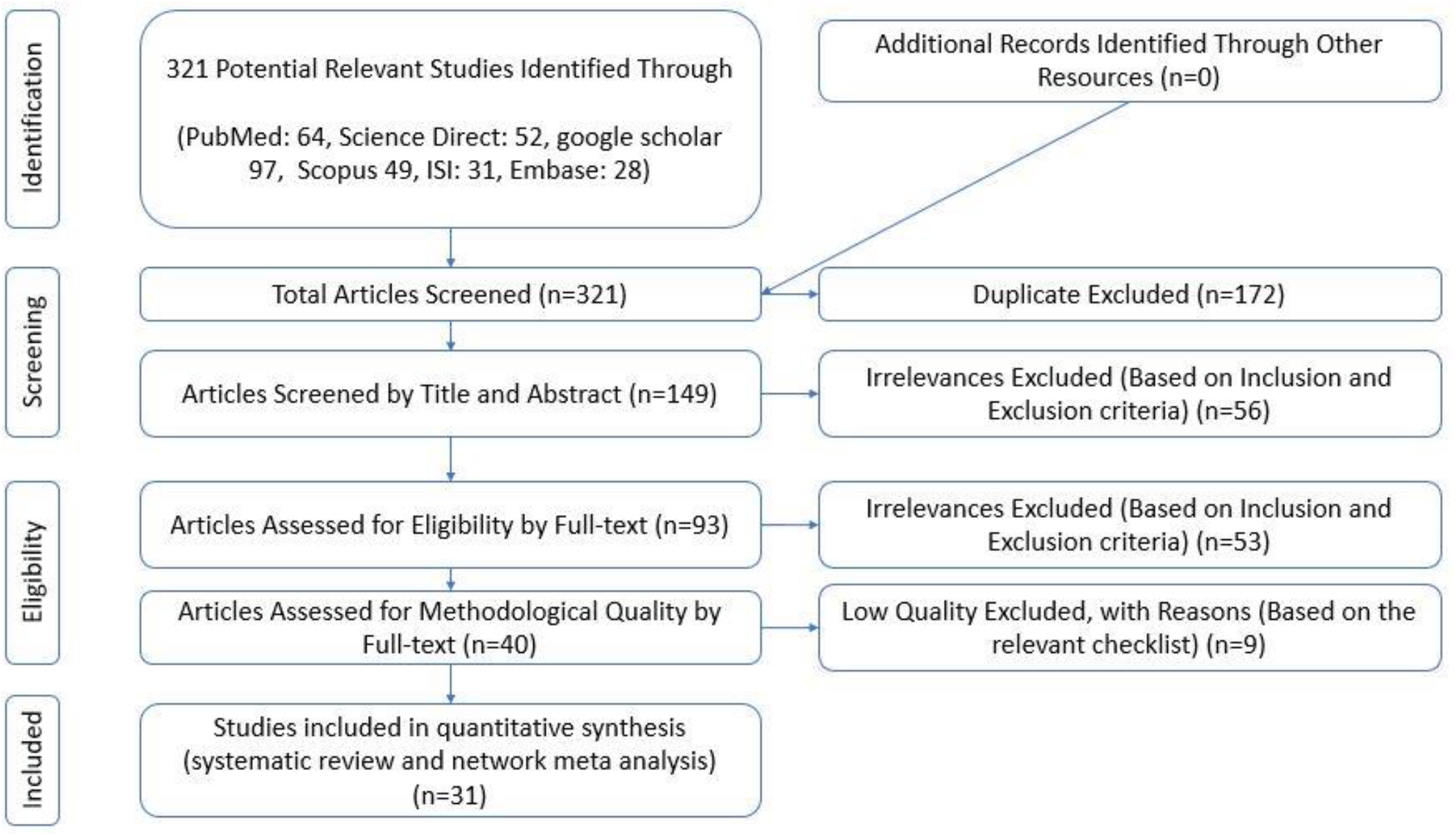
The PRISMA (2020) flowchart illustrates the steps involved in filtering articles within the RAIN protocol.

**Figure 3:**
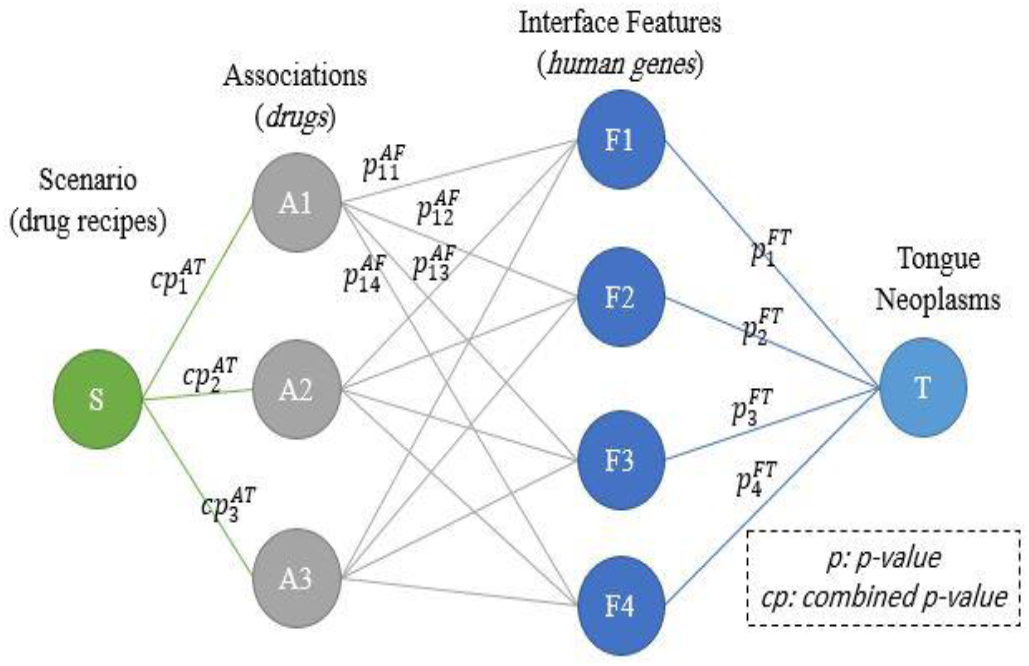
The basic design of the GNN model aims to propose an efficient combination of drugs for treating a disease, utilizing human proteins/genes as interface features.

### 2.2 Stage II: An exhaustive systematic review

The current investigation employed systematic review and meta-analysis techniques following the PRISMA 2020 guidelines. These methods encompassed several stages including identification, screening, eligibility, and inclusion to ensure rigorousness. To mitigate potential errors and publication bias, all phases, including searching, evaluation, identification, selection of articles, and data extraction, were independently conducted by two researchers. Any disparities were resolved through consultation with a supervisor to reach consensus.

#### Study Identification

To locate pertinent records, a comprehensive search was conducted across Persian databases (SID and MagIran) and international ones (PubMed, Embase, Scopus, and Web of Science). The search strategy in each database utilized validated keywords and Medical Subject Headings (MeSH) for PubMed and Emtree for Embase. The search was not restricted by language or time frame, encompassing articles until February 2023. Additionally, manual searches on Google Scholar and reviewing reference lists supplemented the search process.

#### Inclusion Criteria

The inclusion criteria comprised original research articles, observational studies (cross-sectional, cohort, case-control), randomized clinical trials, access to full-text articles, and studies reporting bleeding percentages or frequencies post-PCI.

#### Exclusion Criteria

Qualitative studies, case series, case reports, articles with unavailable full-text after multiple attempts, and redundant studies identified across different databases were excluded.

#### Study Selection Process

Unique search strategies were tailored for each database, and duplicates were removed initially. Author, institution, and journal information were anonymized. Titles and abstracts underwent scrutiny, followed by full-text review against inclusion and exclusion criteria. Articles meeting the criteria underwent qualitative assessment.

#### Quality Assessment

Observational studies were assessed using the STROBE checklist, with scores ≥16 indicating good to moderate methodological quality, while scores <16 resulted in exclusion.

#### Data Extraction

##### Statistical Analysis

Two researchers extracted data utilizing a pre-established checklist encompassing various parameters.

Data were analyzed using Comprehensive Meta-Analysis Software Version 2. Heterogeneity was evaluated using the I2 test, and publication bias was assessed through the Egger test and funnel plot.

The systematic review and meta-analysis adhered to PRISMA guidelines, encompassing 1879 articles from diverse databases. Following removal of duplicates and screening, eight studies met the inclusion criteria after quality assessment.^73^

### 2.3 Stage III: Network meta analysis

Network Meta-Analysis (NMA) is a statistical approach enabling the simultaneous comparison of multiple treatments within a single analysis. This method proves particularly valuable in pharmacogenomics, where understanding how drugs perform concerning specific human genes is pivotal.

#### Data Collection and Selection

In this study, we systematically combed through databases for randomized controlled trials (RCTs) and observational studies detailing the effectiveness of drugs targeting specific human genes associated with [Disease Name]. Our search strategy aimed to encompass studies offering direct or indirect evidence on the relationship between drugs, genes, and the disease.

#### Statistical Procedure

The NMA was executed employing a Bayesian framework, facilitating the integration of prior knowledge and the derivation of probability distributions for each parameter of interest. We established a network of comparisons across diverse studies, ensuring a common comparator was present to interlink disparate data sources.

#### Model Development

We employed a hierarchical model accommodating the multi-level structure of the data, with genes at the primary level, drugs at the secondary level, and the overall treatment effect on the disease at the tertiary level. Adjustments were made for potential confounding factors like age, sex, and disease severity.

#### Evaluation of Drug-Gene Relationships

The NMA model furnished estimates of the relative efficacy of each drug concerning the target genes. These estimates were utilized to rank the drugs based on their potential effectiveness in modulating gene expression pertinent to [Disease Name].

#### Identification of Potential Targets

Subsequently, the NMA results were scrutinized to pinpoint potential gene targets for the disease. Genes exhibiting a significant interaction with effective drugs were deemed potential targets for future drug development.

## 3. RESULTS

### 3.1 Stage I: Graph Neural network

The Graph Neural Network (GNN) proposes a drug combination comprising Tretinoin, Asparaginase, and Cytarabine. The importance of this pairing is detailed in Table 1, showcasing the associated p-values. As an illustration, the p-value concerning Tongue Neoplasms and Tretinoin (Scenario 1) is 0.01. However, upon the addition of Asparaginase (Scenario 2), this value notably decreases to 0.000088. Furthermore, in the third scenario, the p-value decreases even further to 3E-5, indicating the positive impact of the effect of this medication combination on controlling the illness. Table 2 offers an understanding of how the p-values fluctuate between Tongue Neoplasms and human proteins/genes under various scenarios. The ‘S0’ column displays the p-values between Tongue Neoplasms and the corresponding impacted human proteins/genes.

**Table 1:** p-value between scenarios and Tongue Neoplasms.

| Scenario | Drug Combinations | p - Value |
| --- | --- | --- |
| **S1** | Cisplatin | 1.39E-01 |
| **S2** | S1 + Bleomycin | 1.21E-01 |
| **S3** | S2 + Fluorouracil | 1.16E-01 |

**Table 2:** P-values representing the correlation between Tongue Neoplasms and human proteins/genes following the implementation of various scenarios.

| Association Name | S0 | S1 | S2 | S3 | Association Name | S0 | S1 | S2 | S3 |
| --- | --- | --- | --- | --- | --- | --- | --- | --- | --- |
| **FAM168A** | 7.18E-05 | 1.00 | 1.00 | 1.00 | **CASC18** | 1.29E-03 | 0.00 | 0.00 | 0.00 |
| **KRT16P6** | 3.23E-04 | 0.00 | 0.00 | 0.00 | **VIM** | 1.35E-03 | 1.00 | 1.00 | 1.00 |
| **FNDC1-IT1** | 3.23E-04 | 0.00 | 0.00 | 0.00 | **SPRR2G** | 1.62E-03 | 0.00 | 0.00 | 0.00 |
| **MAU2** | 3.64E-04 | 0.99 | 0.99 | 1.00 | **MANBAL** | 1.62E-03 | 0.00 | 0.00 | 0.00 |
| **ASPSCR1** | 5.32E-04 | 0.00 | 0.00 | 0.00 | **C8orf33** | 1.62E-03 | 0.00 | 0.00 | 0.00 |
| **PRAMENP** | 6.46E-04 | 0.00 | 0.00 | 0.00 | **C9orf84** | 1.62E-03 | 0.00 | 0.00 | 0.00 |
| **LINC01527** | 6.46E-04 | 0.99 | 0.99 | 1.00 | **MTFR1** | 1.79E-03 | 0.00 | 0.00 | 0.00 |
| **CDKN2A** | 7.52E-04 | 1.00 | 1.00 | 1.00 | **RAB6C** | 1.79E-03 | 0.00 | 0.00 | 0.00 |
| **CSNK1D** | 8.39E-04 | 0.00 | 0.00 | 0.00 | **MMP2** | 1.83E-03 | 1.00 | 1.00 | 1.00 |
| **NUDCD2** | 9.70E-04 | 0.00 | 0.00 | 0.00 | **ZNF471** | 1.95E-03 | 0.00 | 0.00 | 0.00 |
| **FAM25A** | 9.70E-04 | 0.00 | 0.00 | 0.00 | **MMP9** | 2.46E-03 | 0.99 | 1.00 | 1.00 |
| **KRT16P3** | 9.70E-04 | 0.00 | 0.00 | 0.00 | **ORAOV1** | 2.61E-03 | 0.00 | 0.00 | 0.00 |
| **FAM25BP** | 9.70E-04 | 0.00 | 0.00 | 0.00 | **BZX** | 2.66E-03 | 0.85 | 0.85 | 0.99 |
| **FAM25G** | 9.70E-04 | 0.00 | 0.00 | 0.00 | **DES** | 2.79E-03 | 0.00 | 0.69 | 0.95 |
| **LINC00534** | 9.70E-04 | 0.00 | 0.00 | 0.00 | **BATF2** | 2.81E-03 | 0.92 | 1.00 | 1.00 |
| **LINC01525** | 9.70E-04 | 0.99 | 0.99 | 1.00 | **VCPKMT** | 2.92E-03 | 0.00 | 0.00 | 0.00 |
| **LINC01526** | 9.70E-04 | 0.99 | 0.99 | 1.00 | **TWISTNB** | 2.92E-03 | 0.00 | 0.00 | 0.00 |
| **DNAJB7** | 9.94E-04 | 0.97 | 1.00 | 1.00 | **KRT36** | 2.94E-03 | 0.00 | 0.00 | 0.00 |
| **TP53** | 1.07E-03 | 1.00 | 1.00 | 1.00 | **FRMD4A** | 2.94E-03 | 0.00 | 0.00 | 0.00 |
| **KRT13** | 1.13E-03 | 0.74 | 0.74 | 0.74 | **CCND1** | 2.97E-03 | 1.00 | 1.00 | 1.00 |
| **VEGFC** | 1.16E-03 | 0.93 | 0.98 | 1.00 | **BCL2** | 2.98E-03 | 1.00 | 1.00 | 1.00 |
| **CDH1** | 1.19E-03 | 1.00 | 1.00 | 1.00 | **MTRNR2L12** | 3.07E-03 | 1.00 | 1.00 | 1.00 |
| **CYP4F35P** | 1.29E-03 | 0.00 | 0.00 | 0.00 | **CASC15** | 3.22E-03 | 0.00 | 0.93 | 0.93 |
| **FAM72C** | 1.29E-03 | 0.00 | 0.00 | 0.00 | **COMMD9** | 3.25E-03 | 0.00 | 0.00 | 0.00 |
| **NAALADL2-AS2** | 1.29E-03 | 0.00 | 0.00 | 0.00 | **ECHDC2** | 3.25E-03 | 0.00 | 0.00 | 0.00 |

When Tretinoin is introduced in the S1 column, the aggregated p-values are shown. Particularly, within the ‘S3’ column, the p-values regarding Tongue Neoplasms and human proteins/genes hit 1, suggesting a decrease in the importance of the targeted proteins/genes.

### 3.2 Stage II: A Thorough Systematic Review

This stage involved a comprehensive investigation into how the specified medications impact the treatment of Tongue Neoplasms. Following a meticulous selection process based on PRISMA principles and the RAIN framework, relevant articles were gathered until June 2024. Initially, 321 potentially pertinent articles were identified and arranged using the EndNote reference system. Following the elimination of duplicates (172 articles), the remaining 149 papers were subject to scrutiny based on predetermined inclusion and exclusion criteria during the review of titles and abstracts. Following this, 40 studies were removed during the screening phase. Out of the 120 studies meeting the eligibility criteria, each underwent a thorough evaluation of the full text. Subsequently, 47 studies were discarded due to their alignment with the inclusion and exclusion criteria. During the quality assessment phase, the remaining 16 studies were scrutinized for methodological strength and STROBE checklist scores, resulting in the exclusion of 15 studies due to insufficient quality. Ultimately, 31 cross-sectional studies were incorporated into the final analysis. The complete texts of these articles were meticulously examined and evaluated using the STROBE checklist (specifics outlined in Figure 3). For further details and characteristics, please consult Table 3. Furthermore, the molecular structures of the drugs utilized in the studies are illustrated in Figure 4, and Table 4 offers information concerning the articles included in our analysis of drug combinations. ^74–95^

**Table 3:**
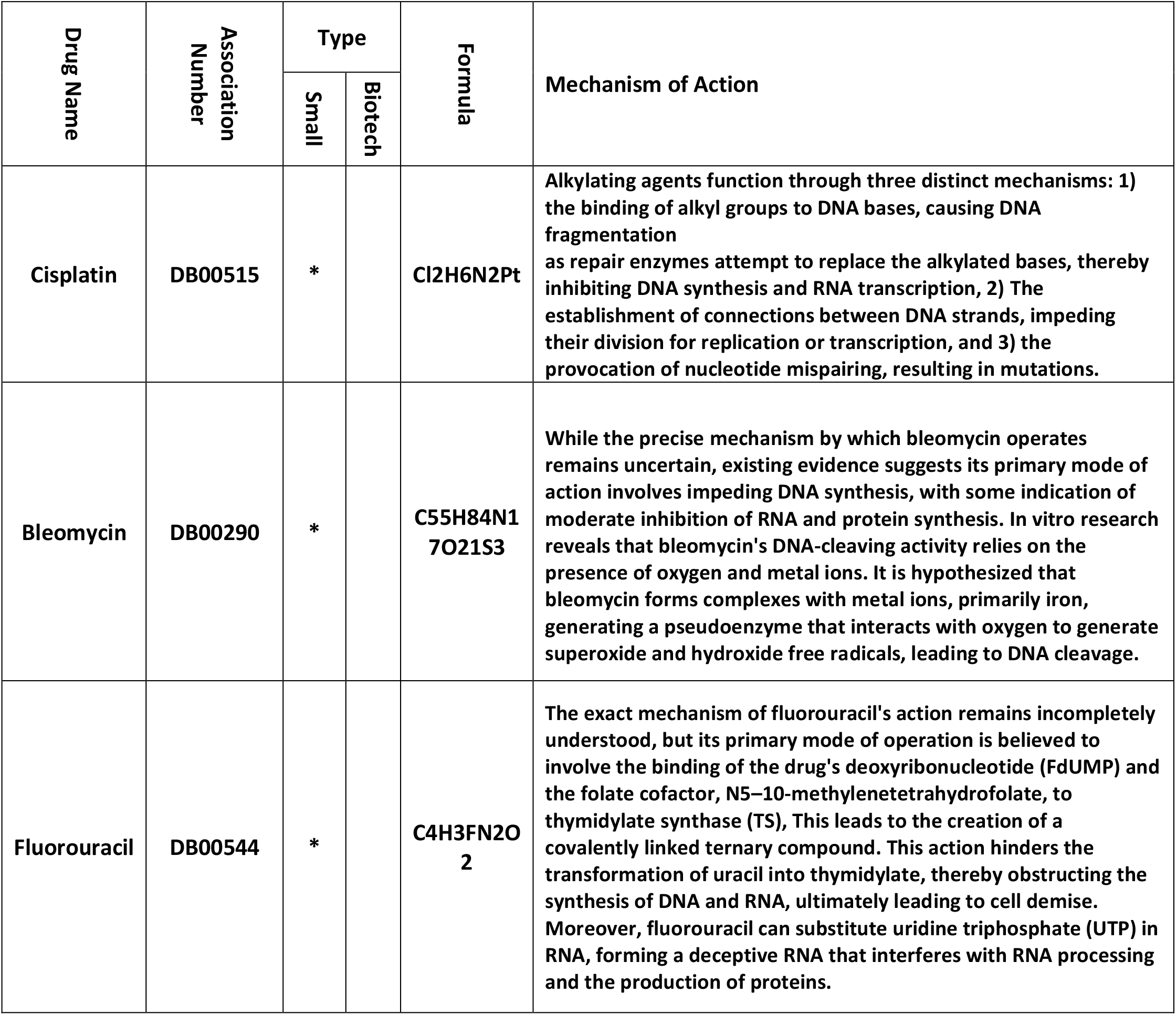
Attributes of recommended drugs as effective therapies for treating Tongue Neoplasms. Name of Drug Accession Code.

| Drug Name | Association Number | Type |  | Formula | Mechanism of Action |
| --- | --- | --- | --- | --- | --- |
|  |  | Small | Biotech |  |  |
| Cisplatin | DB00515 | * |  | Cl<sub>2</sub>H<sub>6</sub>N<sub>2</sub>Pt | Alkylating agents function through three distinct mechanisms: 1) the binding of alkyl groups to DNA bases, causing DNA fragmentation as repair enzymes attempt to replace the alkylated bases, thereby inhibiting DNA synthesis and RNA transcription, 2) The establishment of connections between DNA strands, impeding their division for replication or transcription, and 3) the provocation of nucleotide mispairing, resulting in mutations. |
| Bleomycin | DB00290 | * |  | C<sub>55</sub>H<sub>84</sub>N<sub>17</sub>O<sub>21</sub>S<sub>3</sub> | While the precise mechanism by which bleomycin operates remains uncertain, existing evidence suggests its primary mode of action involves impeding DNA synthesis, with some indication of moderate inhibition of RNA and protein synthesis. In vitro research reveals that bleomycin's DNA-cleaving activity relies on the presence of oxygen and metal ions. It is hypothesized that bleomycin forms complexes with metal ions, primarily iron, generating a pseudoenzyme that interacts with oxygen to generate superoxide and hydroxide free radicals, leading to DNA cleavage. |
| Fluorouracil | DB00544 | * |  | C<sub>4</sub>H<sub>3</sub>FN<sub>2</sub>O<sub>2</sub> | The exact mechanism of fluorouracil's action remains incompletely understood, but its primary mode of operation is believed to involve the binding of the drug's deoxyribonucleotide (FdUMP) and the folate cofactor, N<sup>5</sup>-10-methylenetetrahydrofolate, to thymidylate synthase (TS). This leads to the creation of a covalently linked ternary compound. This action hinders the transformation of uracil into thymidylate, thereby obstructing the synthesis of DNA and RNA, ultimately leading to cell demise. Moreover, fluorouracil can substitute uridine triphosphate (UTP) in RNA, forming a deceptive RNA that interferes with RNA processing and the production of proteins. |

**Table 4:** Some important research studies for proposed drugs in Tongue Neoplasms managements.

| study name | Cisplatin | Bleomycin | Fluorouracil |
| --- | --- | --- | --- |
| Nomura M. et al., 2022 | * |  | * |
| Zhurakivska K. et al., 2022 | * |  | * |
| Hattori M. et al., 2021 | * |  | * |
| Lim SH. et al., 2021 | * |  | * |
| Sakamoto Y. et al., 2019 | * |  | * |
| Kina S. et al., 2019 |  | * | * |
| Huang X. et al., 2018 | * |  | * |
| Tamura T. et al., 2018 | * |  | * |
| Hirabayashi T. et al., 2017 | * |  | * |
| Stojanović N. et al., 2016 | * |  | * |
| Harada K. et al., 2016 | * |  | * |
| Otowa Y. et al., 2015 | * |  | * |
| Chhatui B. et al., 2015 | * |  | * |
| Gebhardt BJ. et al., 2014 | * |  | * |
| Mouawad F. et al., 2014 | * |  | * |
| Gao W. et al., 2014 | * |  | * |
| Matsumoto K. et al., 2013 | * | * | * |
| Matsumoto K. et al., 2013 | * | * |  |
| Matsumoto K. et al., 2013 |  | * | * |
| Peng B. et al., 2012 | * | * | * |
| Peng B. et al., 2012 | * | * |  |
| Peng B. et al., 2012 |  | * | * |
| Zheng G. et al., 2010 | * | * |  |
| Liu S. et al., 2003 | * | * | * |
| Liu S. et al., 2003 | * | * |  |
| Zhang P. et al., 2003 | * | * |  |
| Zhang P. et al., 2003 | * | * |  |
| Liu S. et al., 2003 |  | * | * |
| Song M. et al., 2002 | * | * | * |
| Song M. et al., 2002 | * | * |  |
| Song M. et al., 2002 |  | * | * |

**Figure 4:**
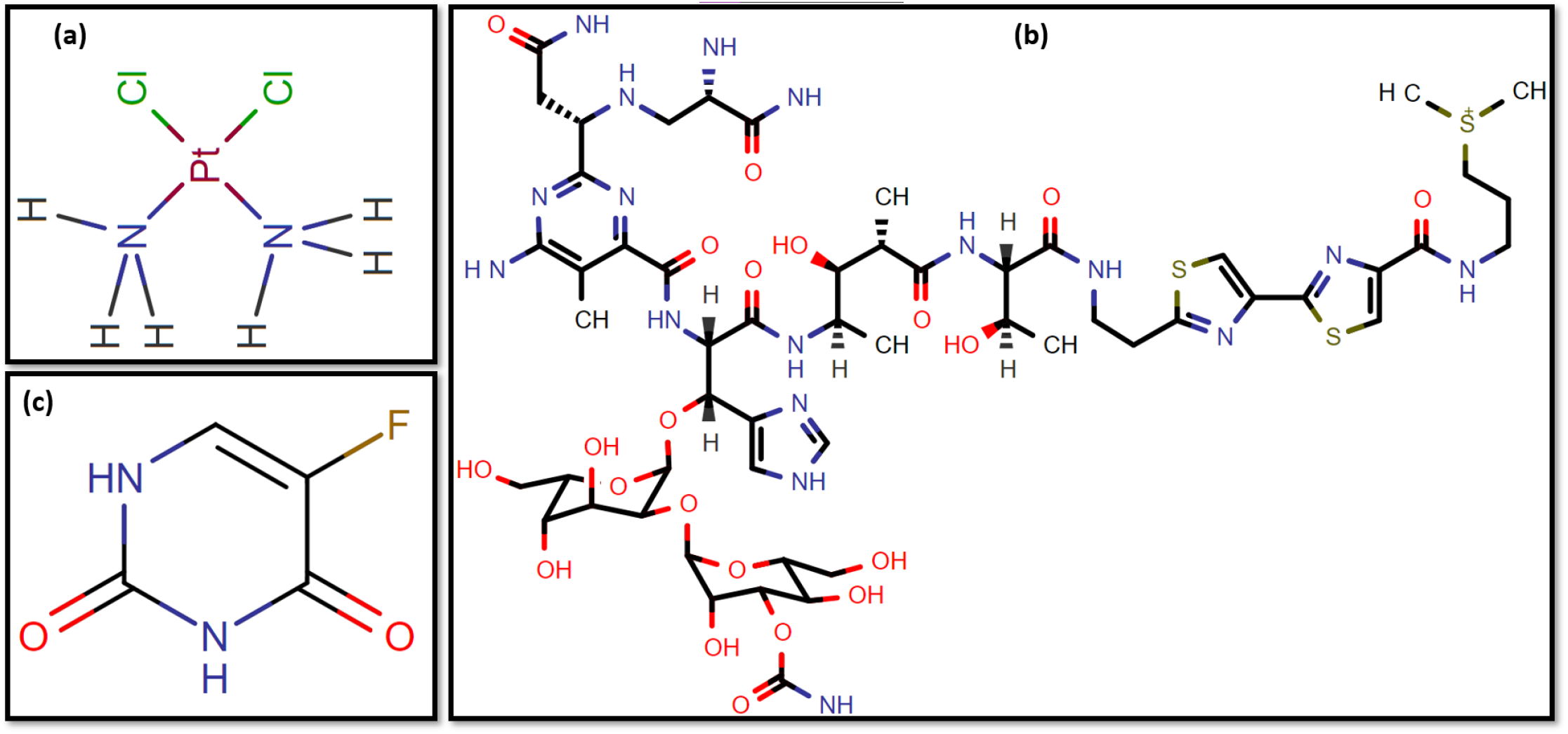
Drug structure for (a) Cisplatin, (b) Bleomycin, (c) Fluorouracil. From https://www.drugbank.com/

### 3.3 Stage III: Network Meta-Analysis

Figure 5 exhibits the p-values linked to human proteins/genes impacted by Tongue Neoplasms, whereas Figure 6 demonstrates the p-values derived from the third scenario. The efficacy of drugs identified by the drug selection algorithm is depicted in Figure 7 using a radar chart, demonstrating the p-values between Tongue Neoplasms and human proteins/genes after the administration of the chosen medications. Figure 8 displays the p-values indicating associations and targets using various interface features. P-values below .01 and .05 are represented in green and blue, respectively. Each colored line indicates the effectiveness of the respective drug in the given scenario.

**Figure 5:**
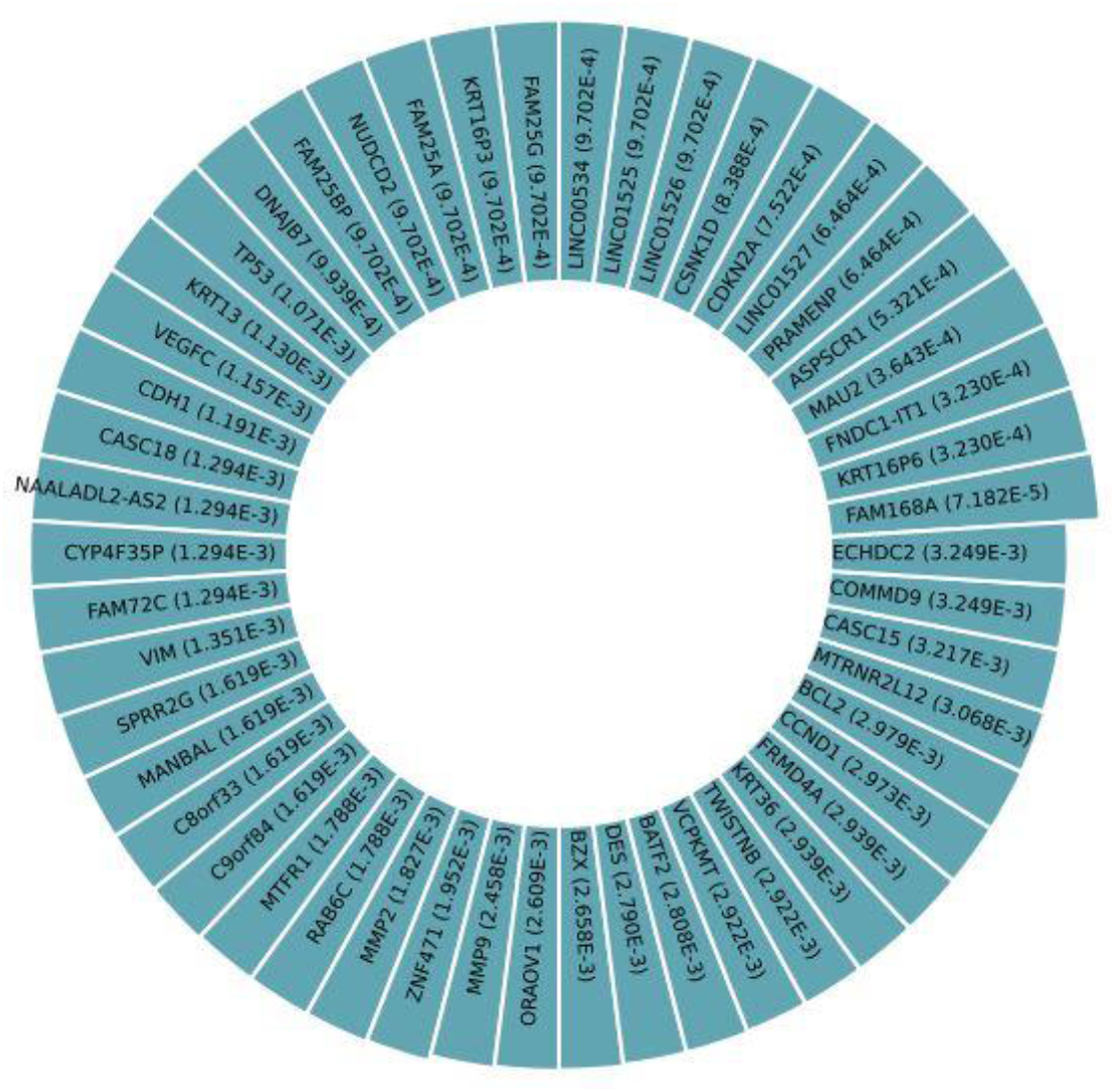
P-values indicating the relationship between impacted human proteins/genes and Tongue Neoplasms.

**Figure 6:**
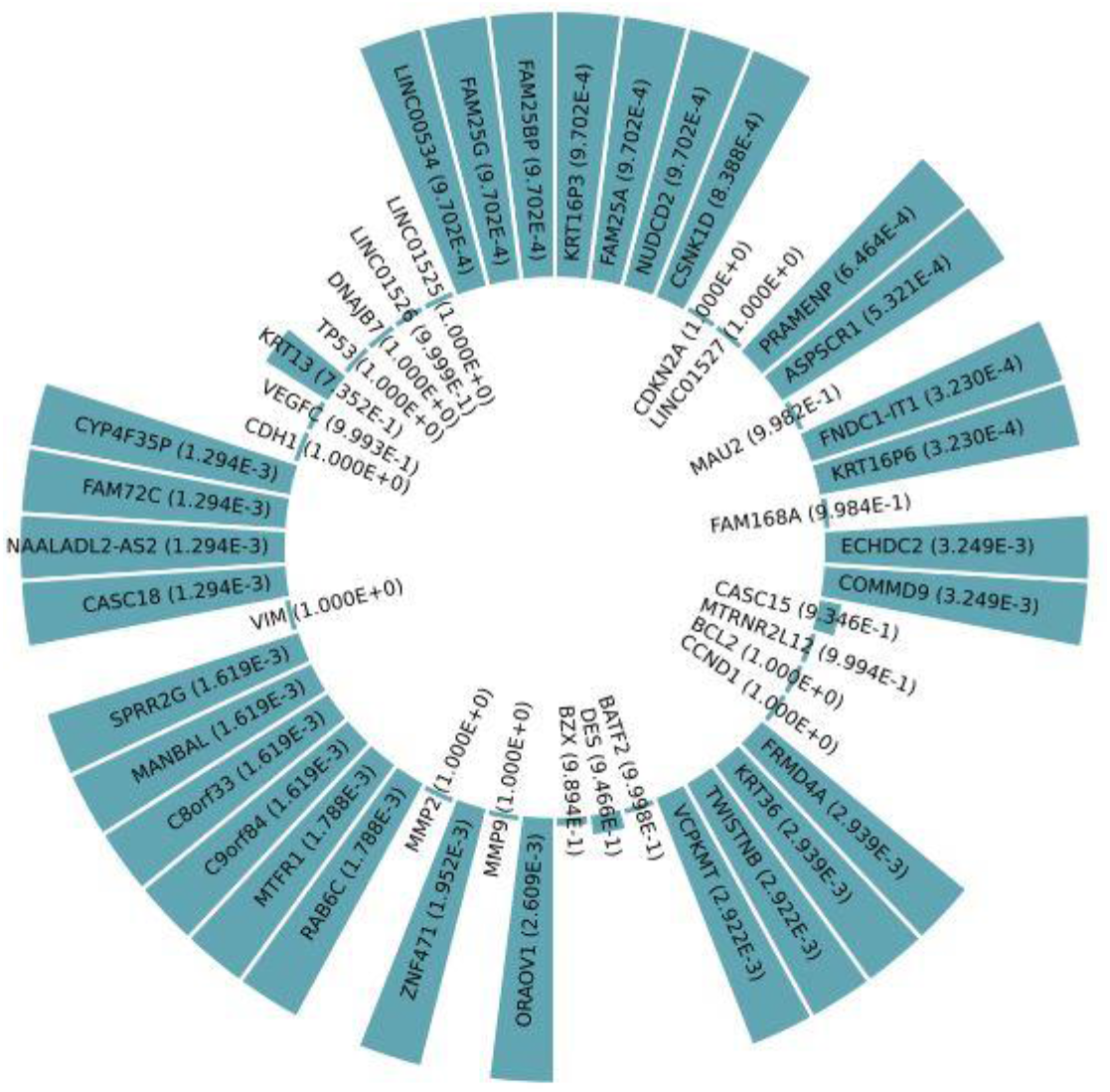
P-values representing the correlation between impacted human proteins/genes and Tongue Neoplasms following the implementation of Scenario 4

**Figure 7:**
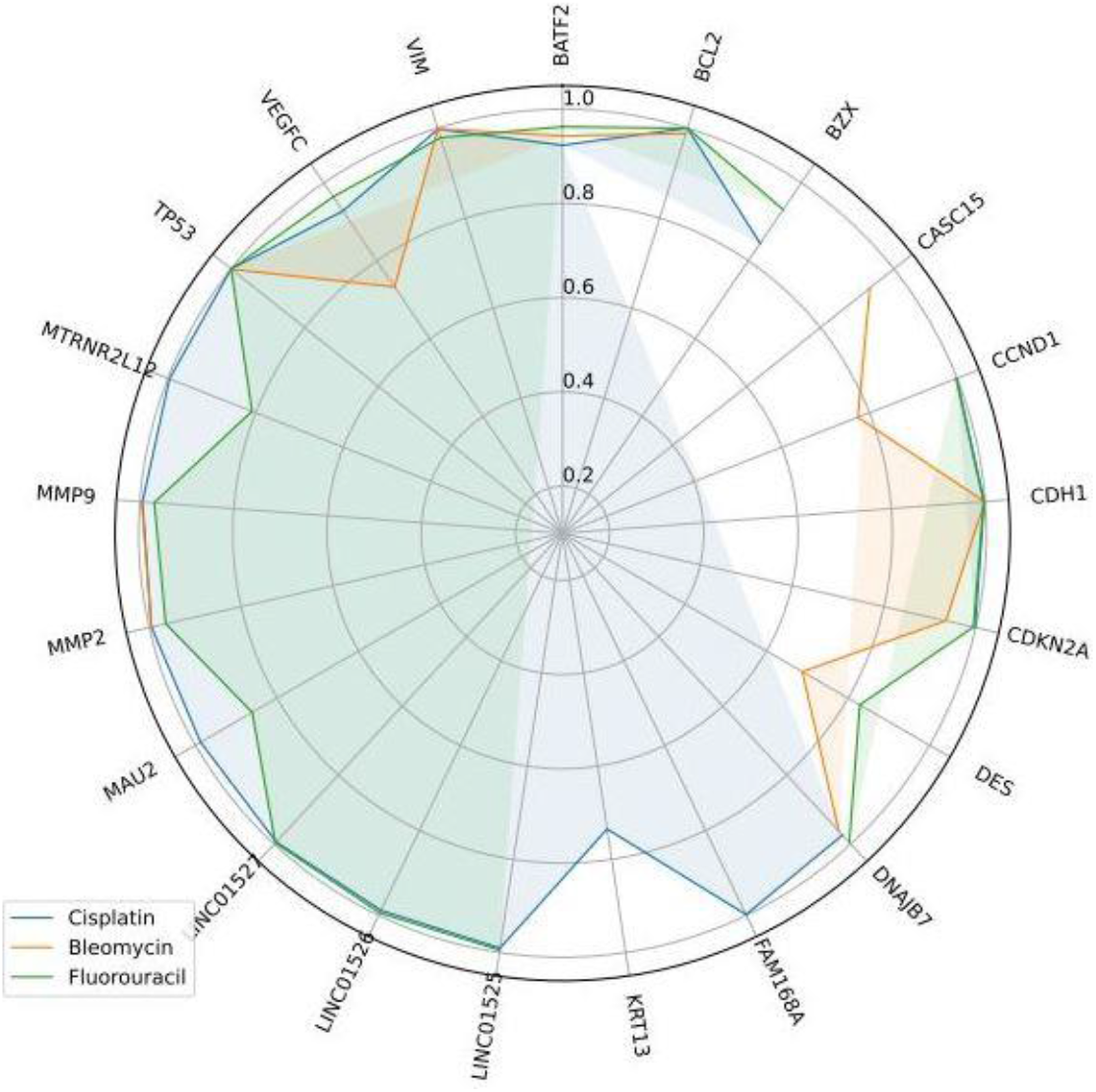
Radar chart displaying the p-values correlating Tongue Neoplasms with affected proteins/genes subsequent to the intake of individual drugs.

**Figure 8:**
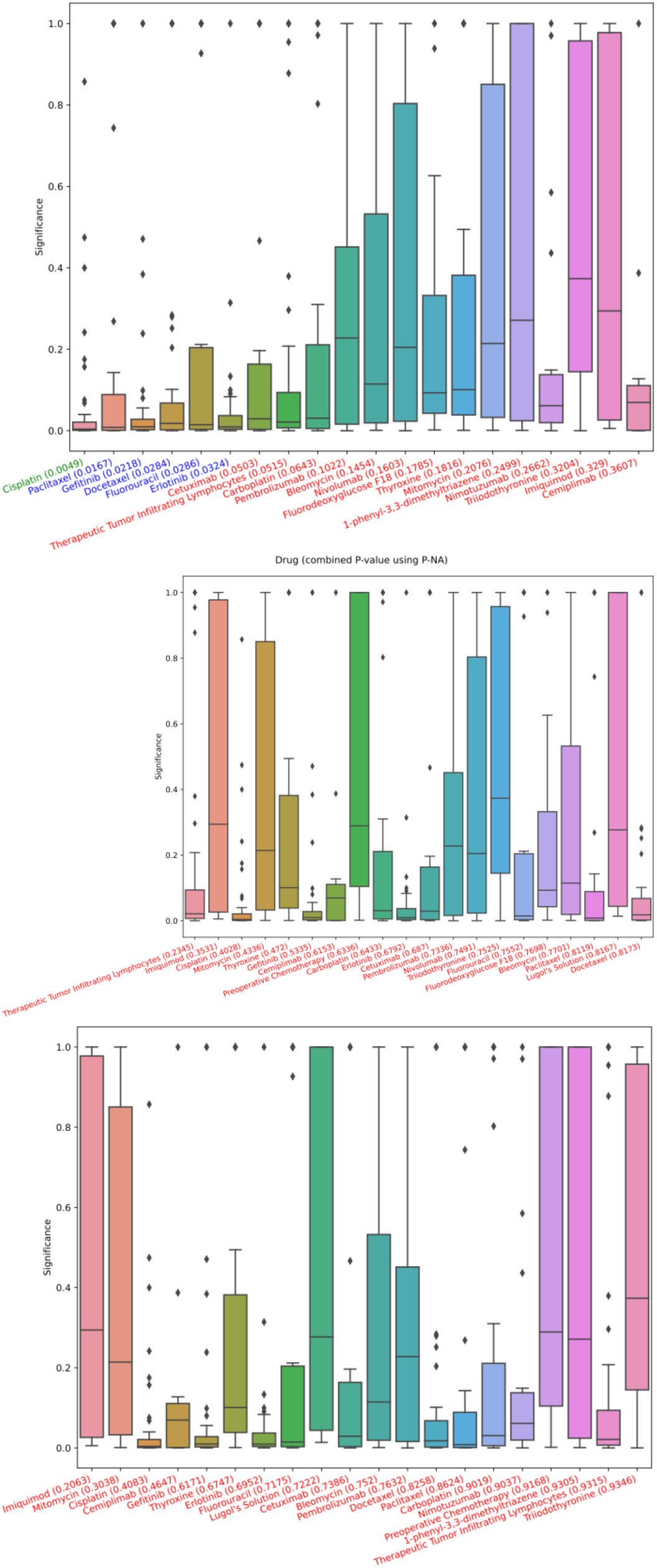
p-values between associations and target, using different interface features. (Top) Overall, (Middle) after first drug from Scenario one is used, (Bottom) after both drugs from Scenario two are used.

## 4. DISCUSSION

Accessing details about prescription drugs is crucial for grasping their possible interactions, adverse reactions, and related risks. To thoroughly compare various medications, reputable online references like Medscape, WebMD, Drugs, and Drugbank were consulted. Employing these trusted databases, an extensive analysis of drug combinations was undertaken, uncovering distinct interactions between certain pairs of medications. Notably, the examined websites emphasized an elevated risk or severity of side effects when combining bleomycin with cisplatin, cisplatin with fluorouracil, or bleomycin with fluorouracil.

## 5. CONCLUSION

This paper introduces an innovative approach to identifying effective drug combinations for treating Tongue Neoplasms, a prevalent and fatal cancer type. Our method integrates DMoN and the RAIN protocol, combining Graph Neural Network, natural language processing, and network meta-analysis techniques. Applying this approach to a dataset linking drugs and genes associated with Tongue Neoplasms, we pinpointed Cisplatin, Bleomycin, and Fluorouracil as promising drugs for targeting the implicated genes/proteins. Validation through literature review and statistical analysis supports the robustness of our findings. Our method provides a new and powerful tool to assist healthcare providers and researchers in customizing optimal treatments for patients with Tongue Neoplasms and understanding the underlying mechanisms of the disease. We anticipate its extension to various cancer types and diseases, potentially advancing precision medicine and drug discovery.

## Declarations

## Ethical Approval and Participation Consent

Not required.

## Publication Consent

Not required.

## Data and Material Availability

The data generated or analyzed for this study are included within the published article.

## Contributions by Authors

MB and AAK conducted the analysis. MA also interpreted the findings. MA was responsible for drafting the manuscript. All authors have read and given their approval for the final version of the manuscript.

## Acknowledgements

The authors extend their gratitude for the financial backing provided by the Research and Technology Department at Kermanshah University of Medical Sciences.

