## Supplemental Figures for "Emerging Drug Combinations for Targeting Tongue Neoplasms Associated Proteins/Genes: Employing Graph Neural Networks within the RAIN Protocol"

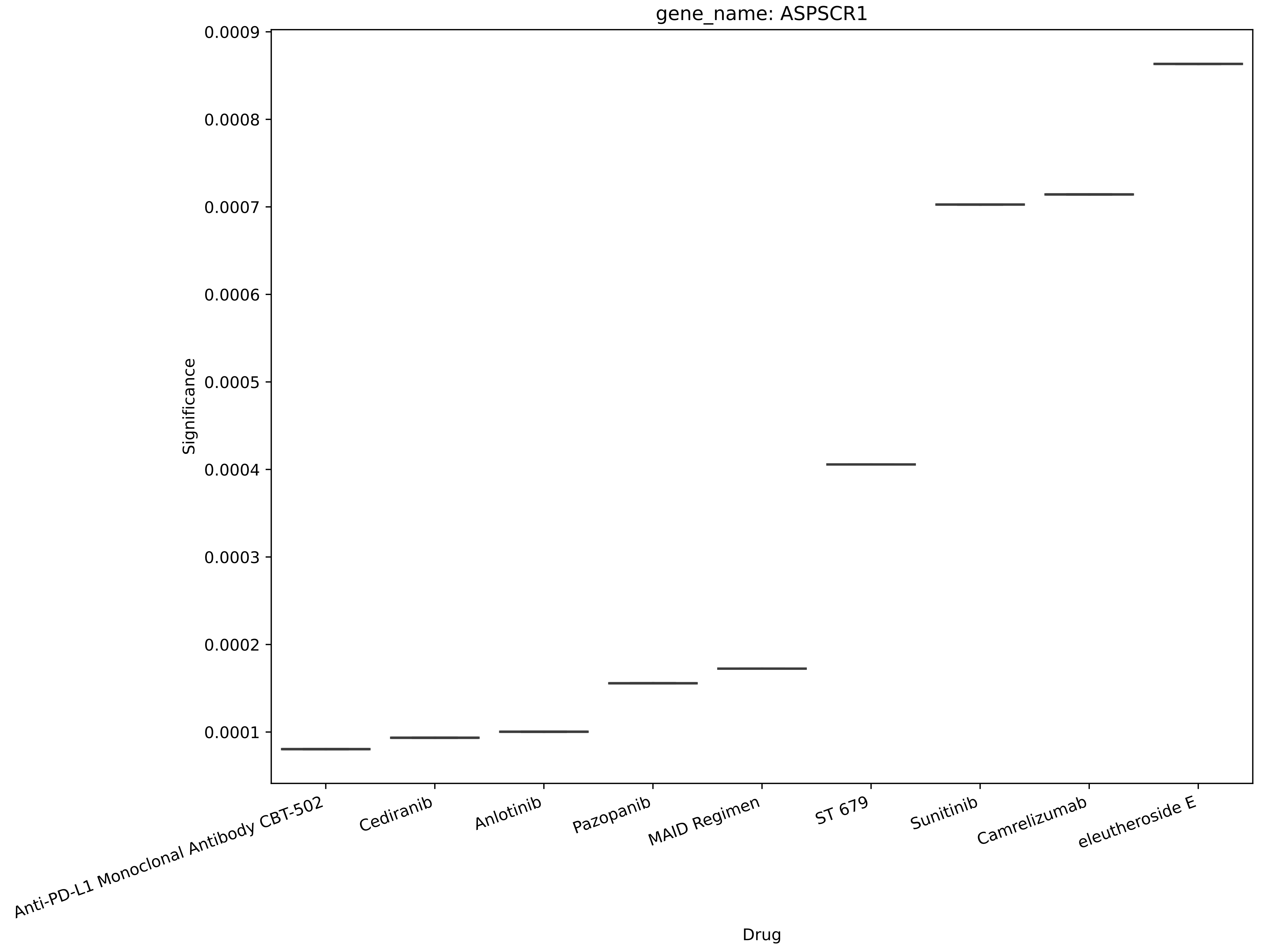
gene_name: ASPSCR1
Significance
0.0009
0.0008
0.0007
0.0006
0.0005
0.0004
0.0003
0.0002
0.0001
Anti-PD-L1 Monoclonal Antibody CBT-502
Cediranib
Anlotinib
Pazopanib
MAID Regimen
ST 679
Sunitinib
Camrelizumab
eleutheroside E
Drug

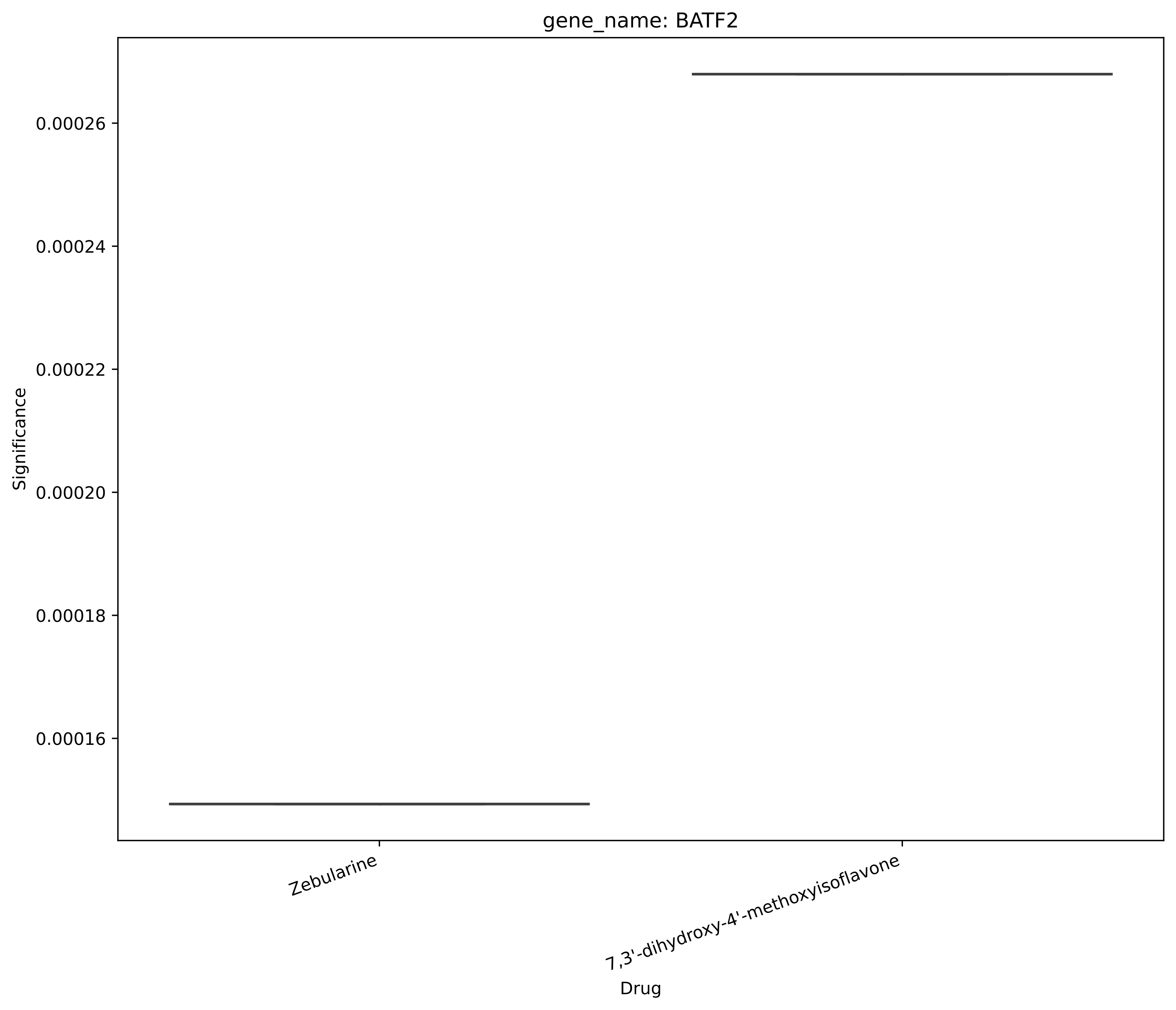
gene_name: BATF2
Significance
0.00026
0.00024
0.00022
0.00020
0.00018
0.00016
Zebularine
7,3'-dihydroxy-4'-methoxyisoflavone
Drug

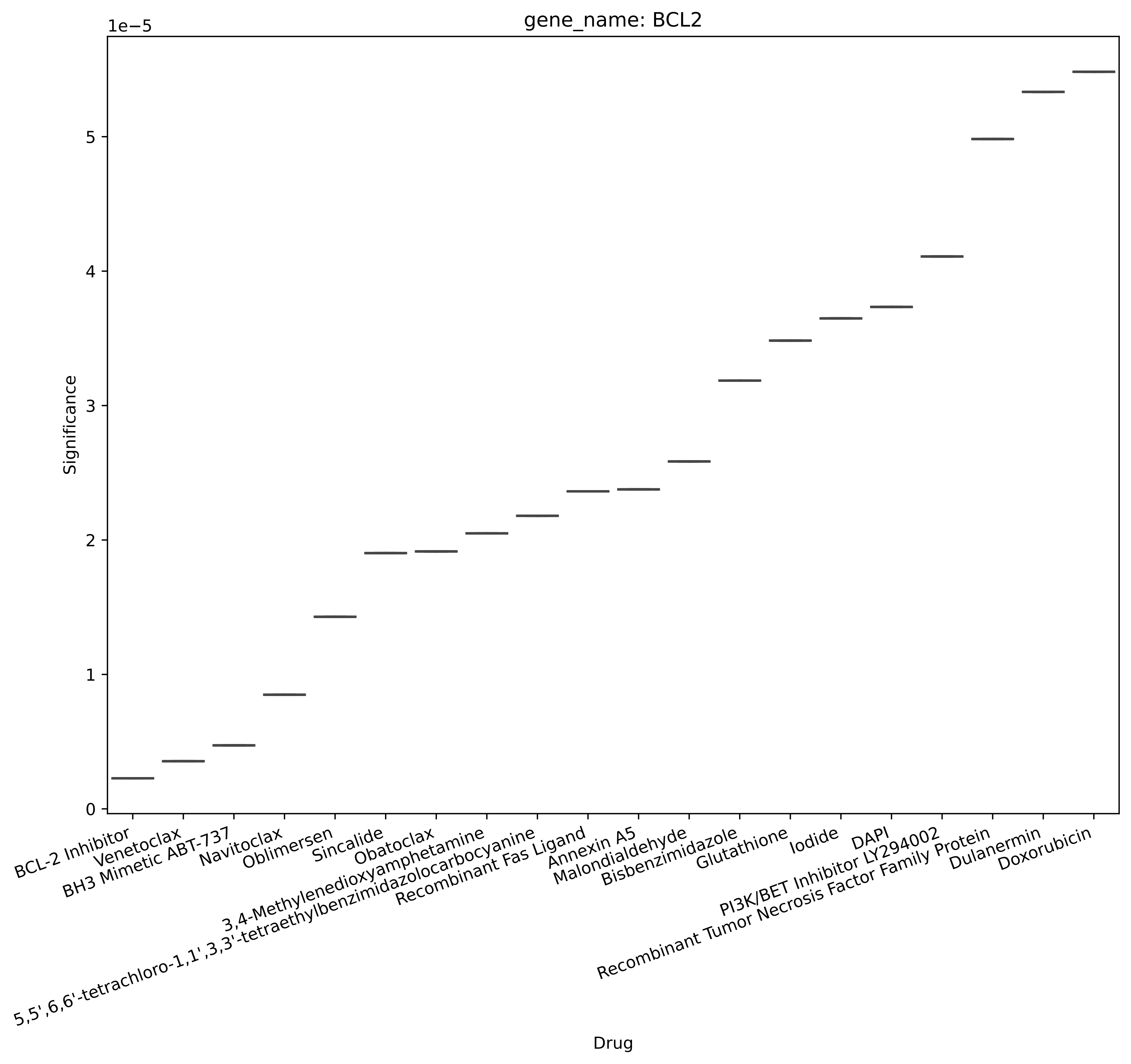
gene_name: BCL2
1e−5
Significance
0
1
2
3
4
5
BCL-2 Inhibitor
Venetoclax
BH3 Mimetic ABT-737
Navitoclax
Oblimersen
Sincalide
Obatoclax
3,4-Methylenedioxyamphetamine
5,5',6,6'-tetrachloro-1,1',3,3'-tetraethylbenzimidazolocarbocyanine
Recombinant Fas Ligand
Annexin A5
Malondialdehyde
Bisbenzimidazole
Glutathione
Iodide
DAPI
PI3K/BET Inhibitor LY294002
Recombinant Tumor Necrosis Factor Family Protein
Dulanermin
Doxorubicin
Drug

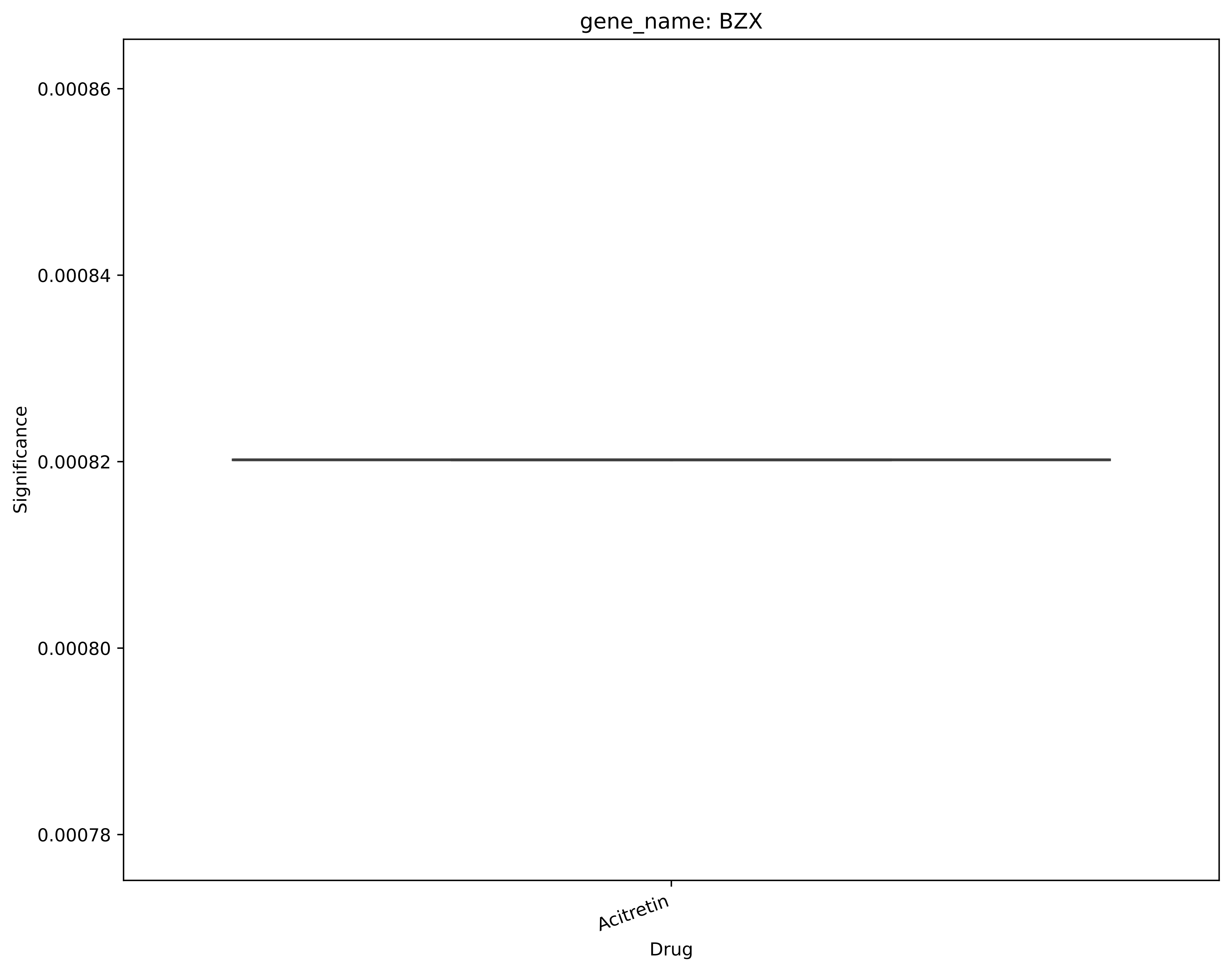
gene_name: BZX
0.00086
0.00084
0.00082
0.00080
0.00078
Significance
Acitretin
Drug

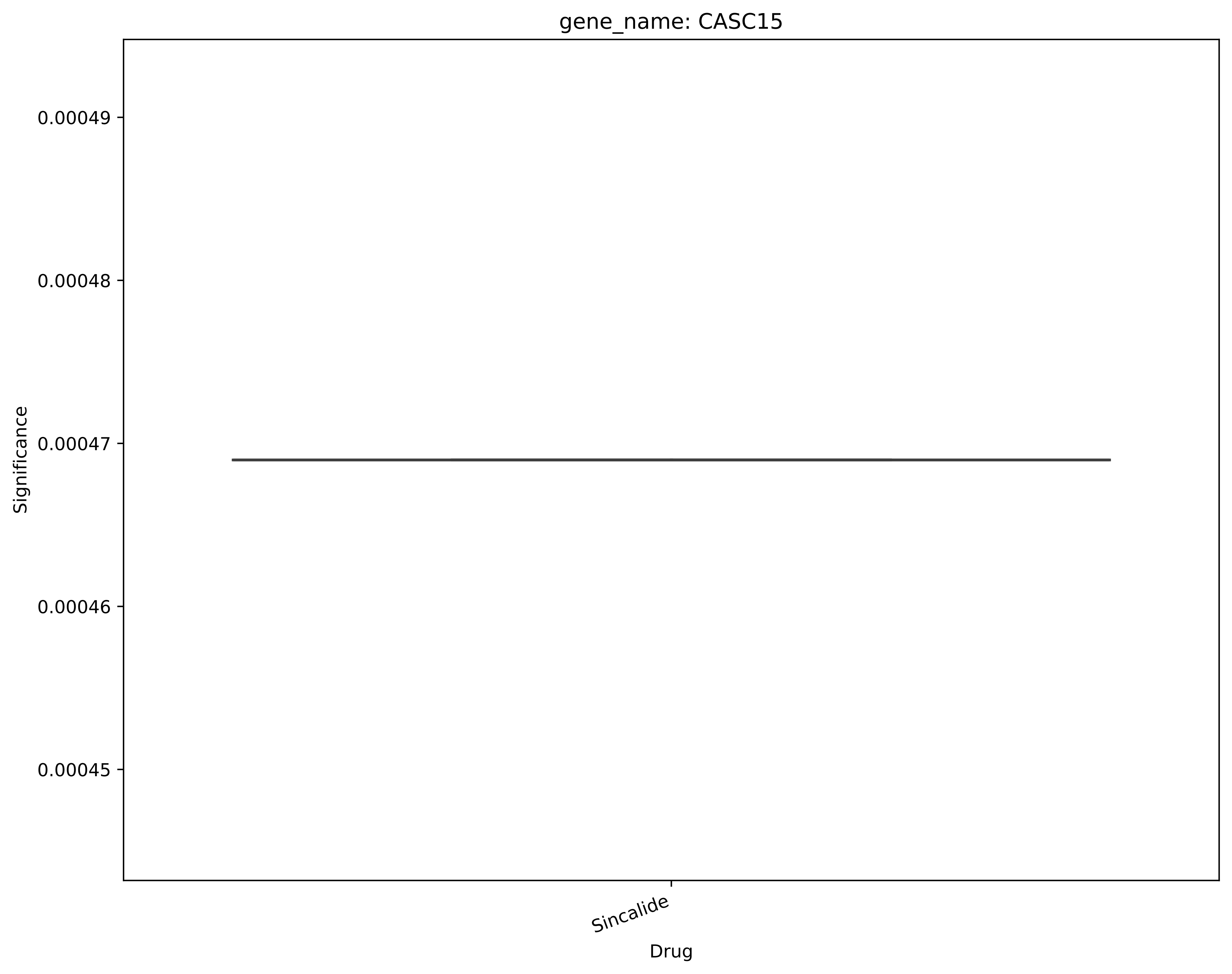
gene_name: CASC15
0.00049
0.00048
0.00047
0.00046
0.00045
Significance
Sincalide
Drug

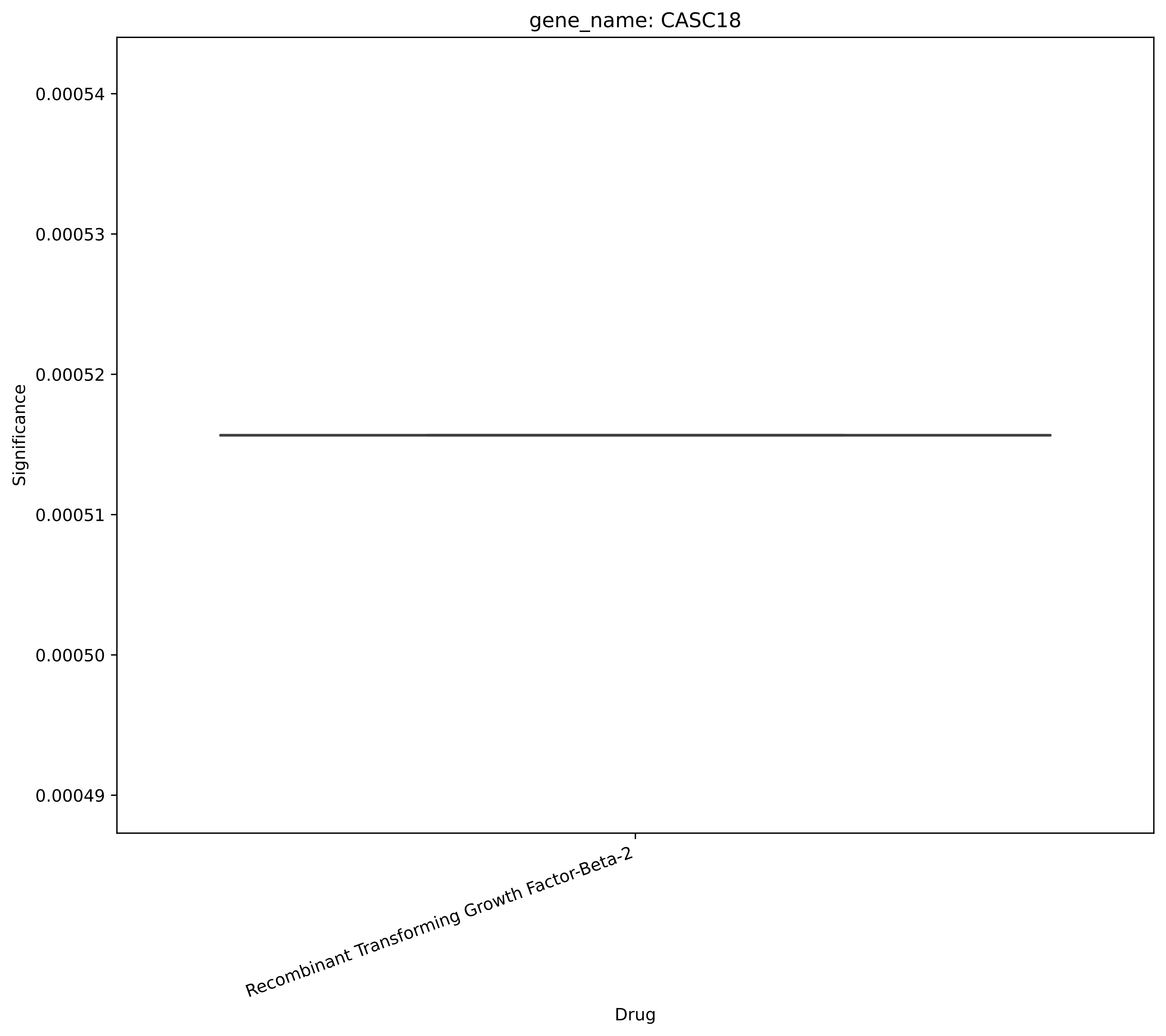

gene_name: CASC18
0.00054
0.00053
0.00052
0.00051
0.00050
0.00049
Significance
Recombinant Transforming Growth Factor-Beta-2
Drug

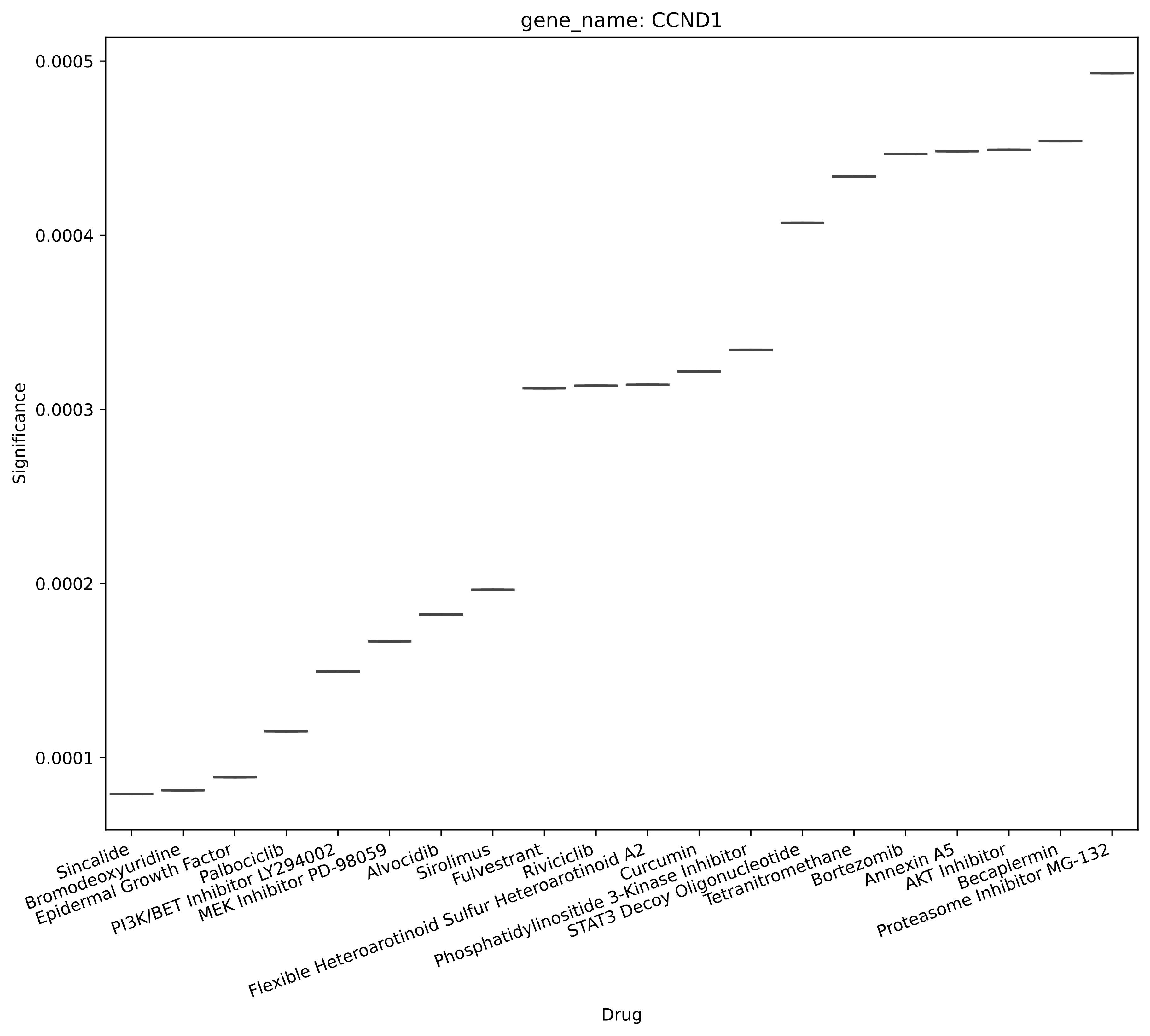
gene_name: CCND1
Significance
0.0005
0.0004
0.0003
0.0002
0.0001
Drug
Sincalide
Bromodeoxyuridine
Epidermal Growth Factor
Palbociclib
PI3K/BET Inhibitor LY294002
MEK Inhibitor PD-98059
Alvocidib
Sirolimus
Fulvestrant
Riviciclib
Flexible Heteroarotinoid Sulfur Heteroarotinoid A2
Curcumin
Phosphatidylinositide 3-Kinase Inhibitor
STAT3 Decoy Oligonucleotide
Tetranitromethane
Bortezomib
Annexin A5
AKT Inhibitor
Becaplermin
Proteasome Inhibitor MG-132

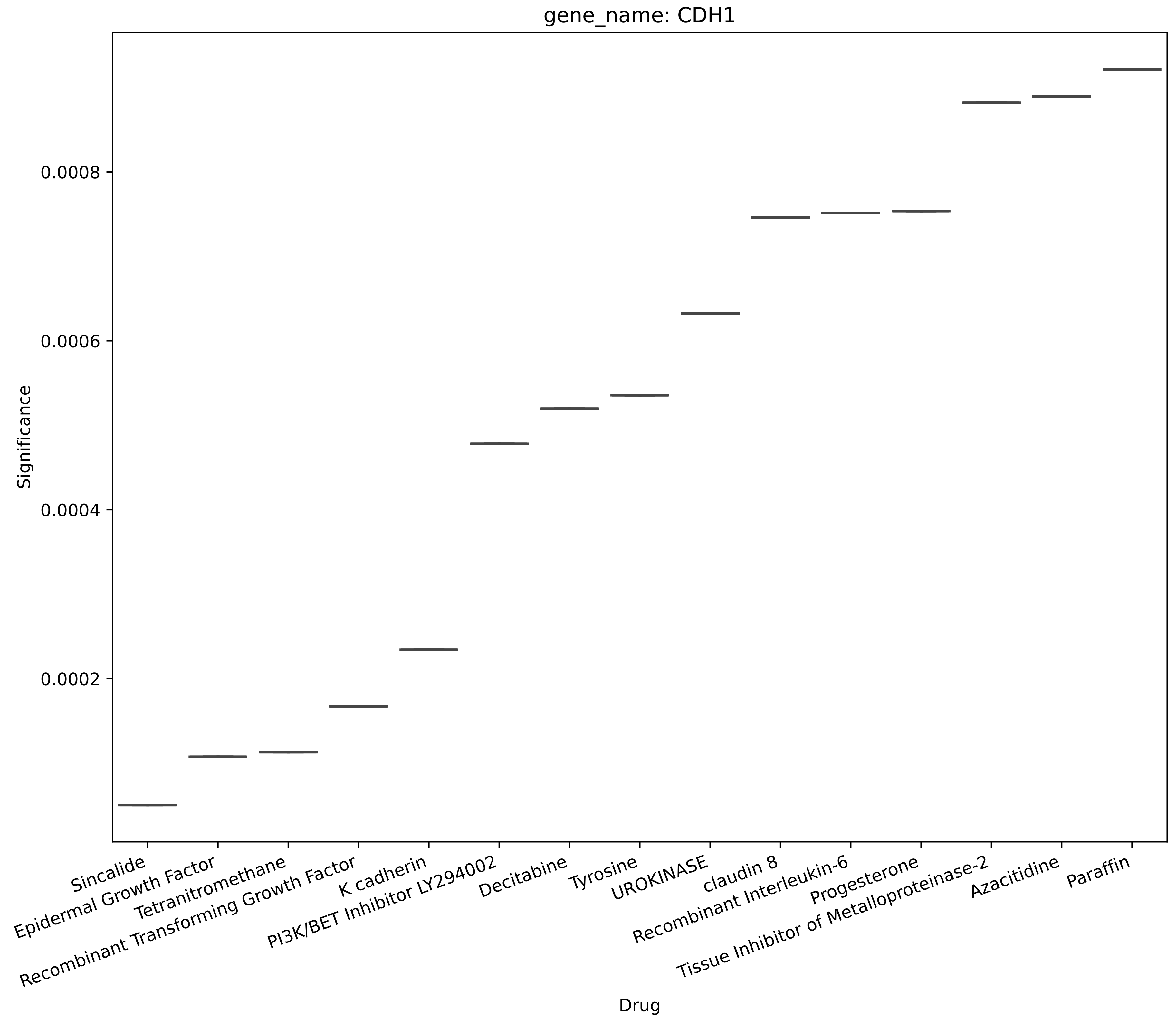
gene_name: CDH1
Significance
0.0008
0.0006
0.0004
0.0002
Drug
Sincalide
Epidermal Growth Factor
Tetranitromethane
Recombinant Transforming Growth Factor
K cadherin
PI3K/BET Inhibitor LY294002
Decitabine
Tyrosine
UROKINASE
claudin 8
Recombinant Interleukin-6
Progesterone
Tissue Inhibitor of Metalloproteinase-2
Azacitidine
Paraffin

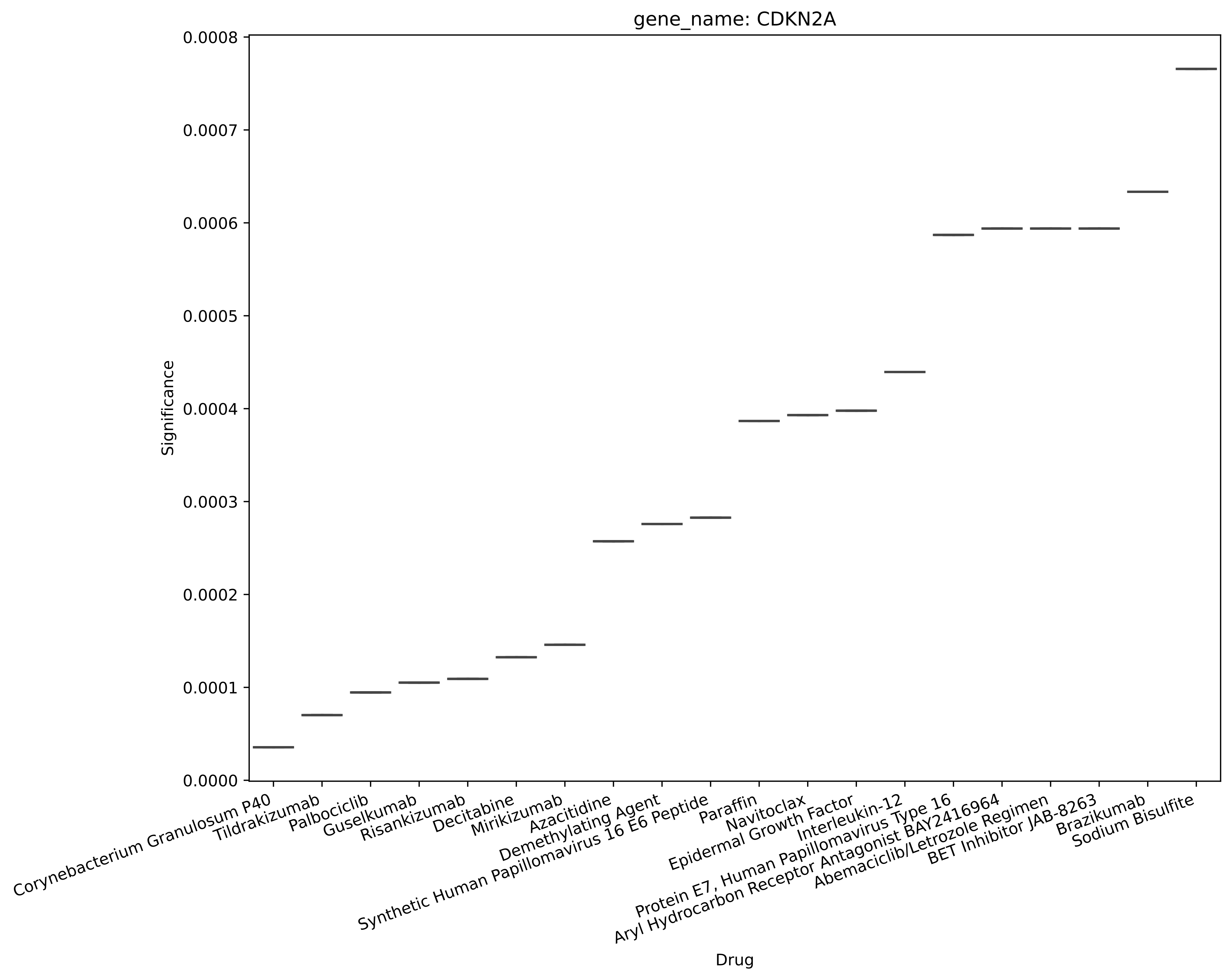
gene_name: CDKN2A
Significance
0.0000
0.0001
0.0002
0.0003
0.0004
0.0005
0.0006
0.0007
0.0008
Corynebacterium Granulosum P40
Tildrakizumab
Palbociclib
Guselkumab
Risankizumab
Decitabine
Mirikizumab
Azacitidine
Demethylating Agent
Synthetic Human Papillomavirus 16 E6 Peptide
Paraffin
Navitoclax
Epidermal Growth Factor
Interleukin-12
Protein E7, Human Papillomavirus Type 16
Aryl Hydrocarbon Receptor Antagonist BAY2416964
Abemaciclib/Letrozole Regimen
BET Inhibitor JAB-8263
Brazikumab
Sodium Bisulfite
Drug

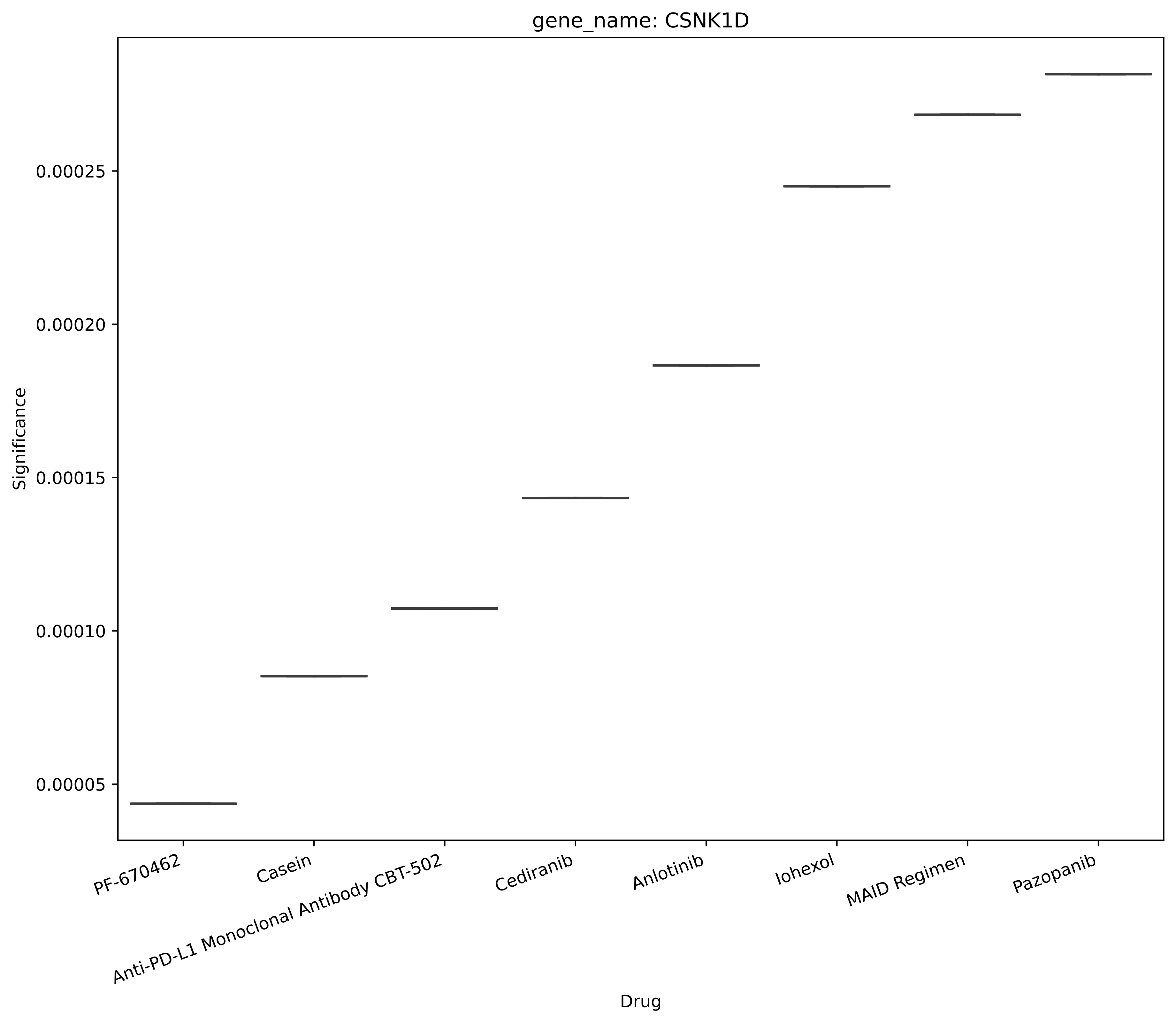
gene_name: CSNK1D
Significance
0.00005
0.00010
0.00015
0.00020
0.00025
PF-670462
Casein
Anti-PD-L1 Monoclonal Antibody CBT-502
Cediranib
Anlotinib
Iohexol
MAID Regimen
Pazopanib
Drug

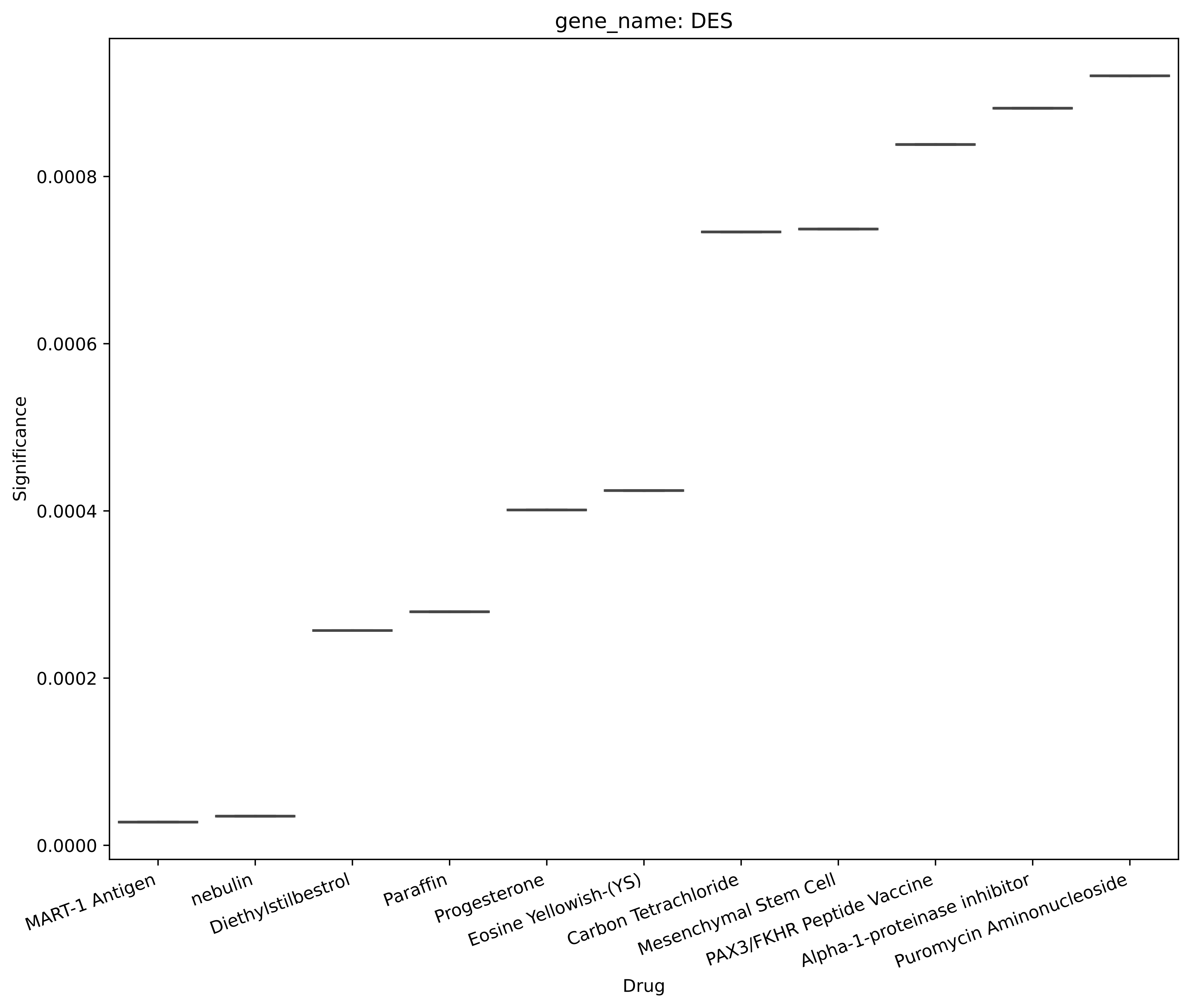

gene_name: DES
Significance
0.0000
0.0002
0.0004
0.0006
0.0008
MART-1 Antigen
nebulin
Diethylstilbestrol
Paraffin
Progesterone
Eosine Yellowish-(YS)
Carbon Tetrachloride
Mesenchymal Stem Cell
PAX3/FKHR Peptide Vaccine
Alpha-1-proteinase inhibitor
Puromycin Aminonucleoside
Drug

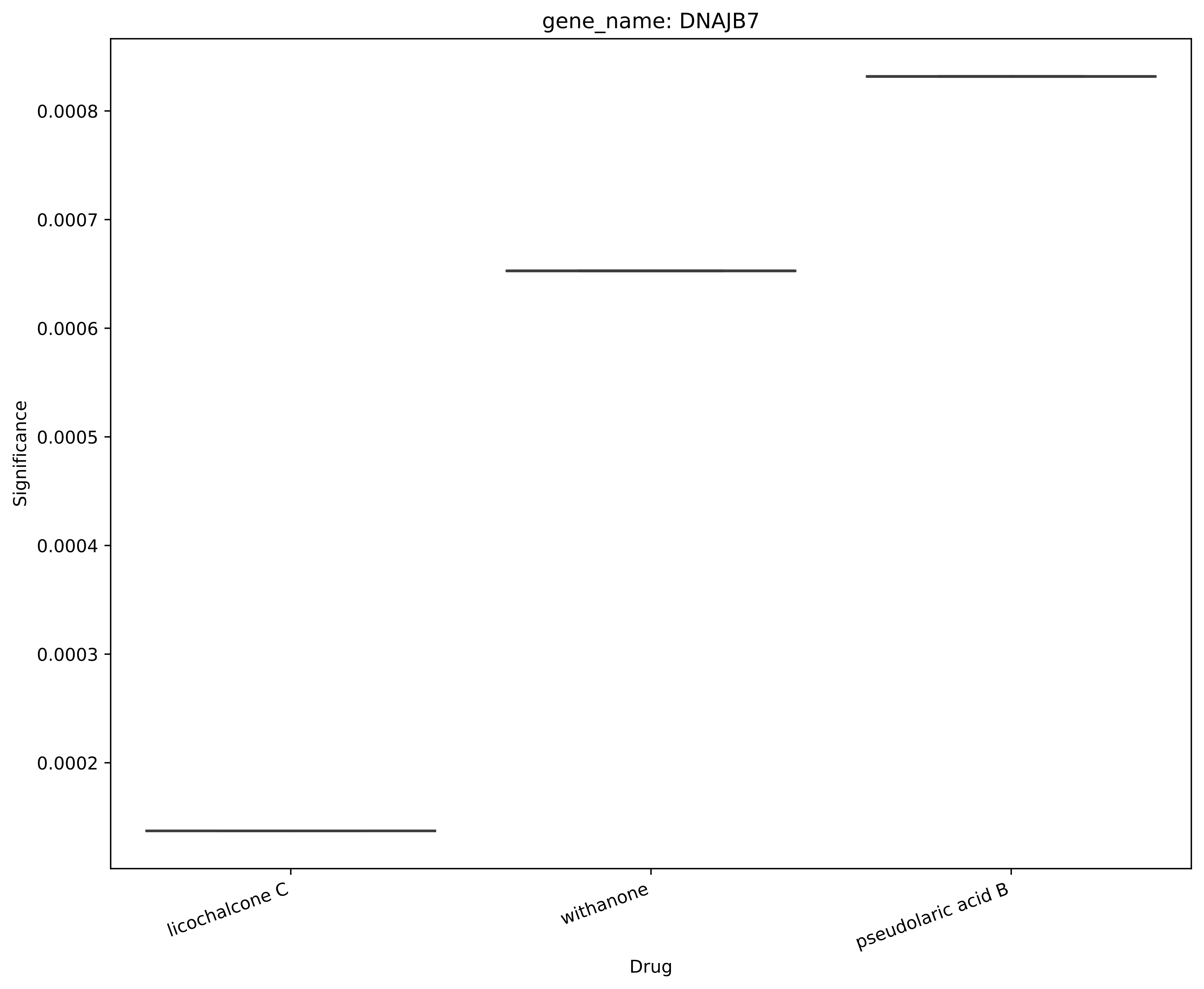
gene_name: DNAJB7
Significance
0.0008
0.0007
0.0006
0.0005
0.0004
0.0003
0.0002
licochalcone C
withanone
pseudolaric acid B
Drug

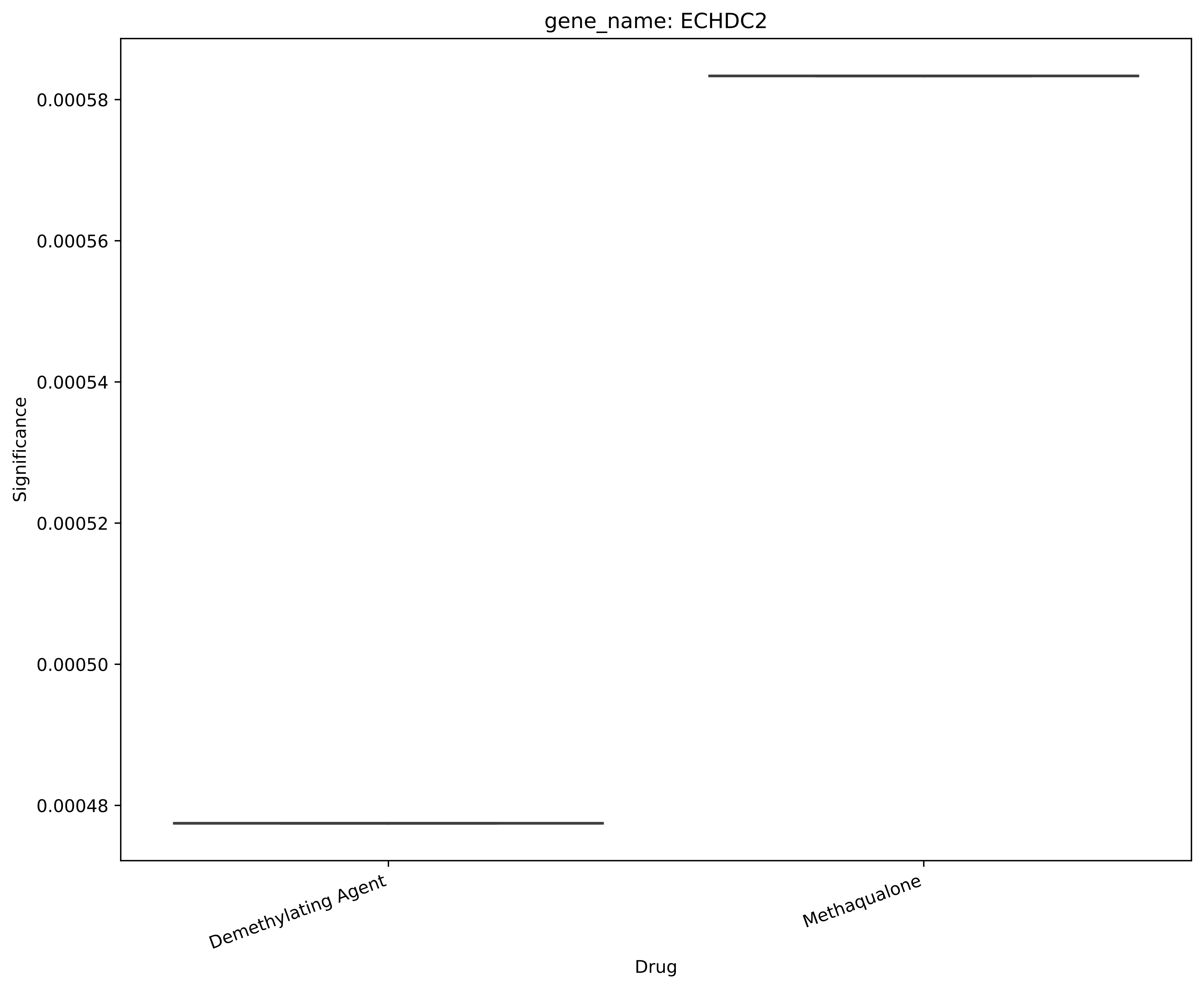
gene_name: ECHDC2
Significance
0.00058
0.00056
0.00054
0.00052
0.00050
0.00048
Demethylating Agent
Methaqualone
Drug

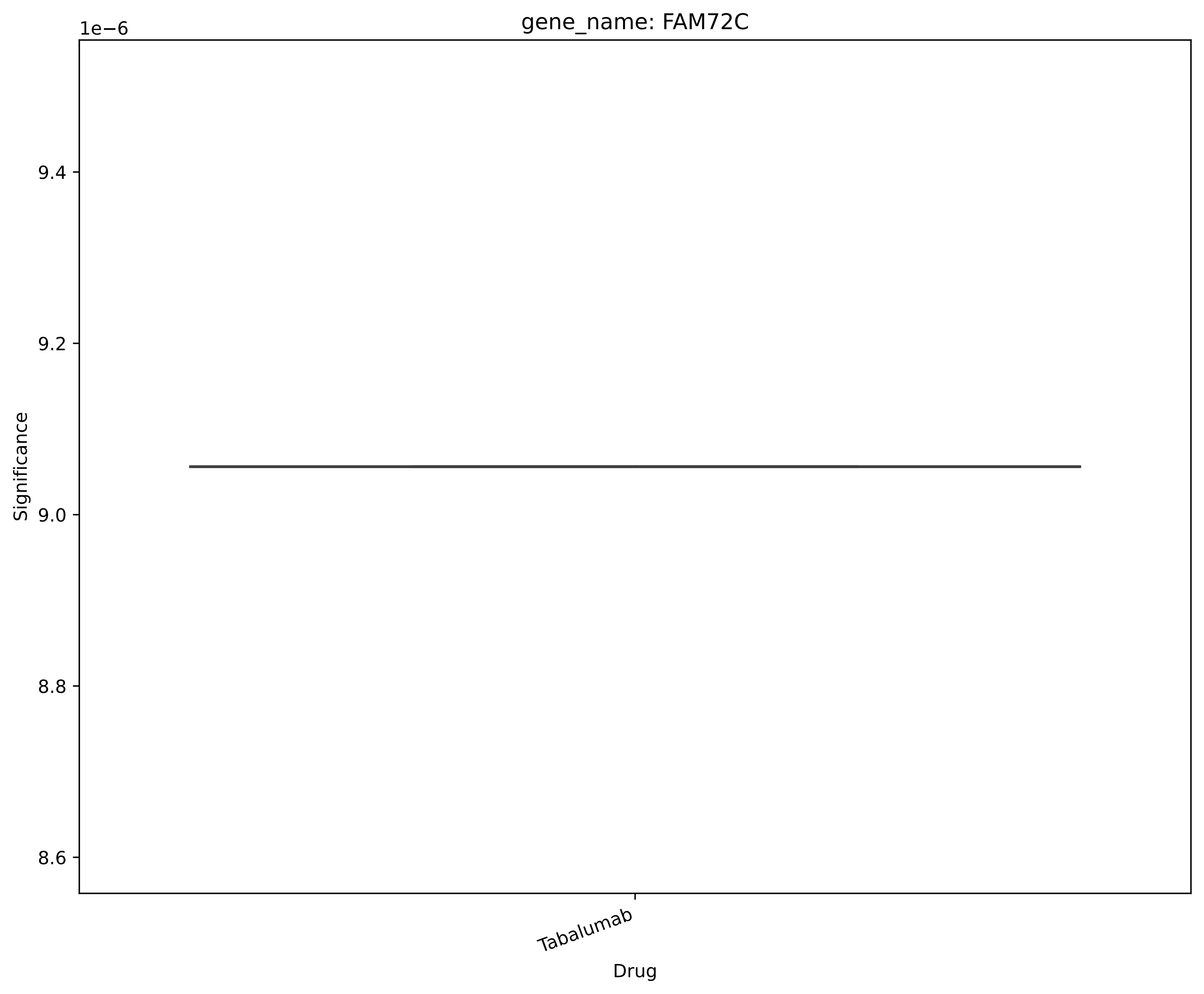
gene_name: FAM72C
1e−6
9.4
9.2
9.0
8.8
8.6
Significance
Tabalumab
Drug

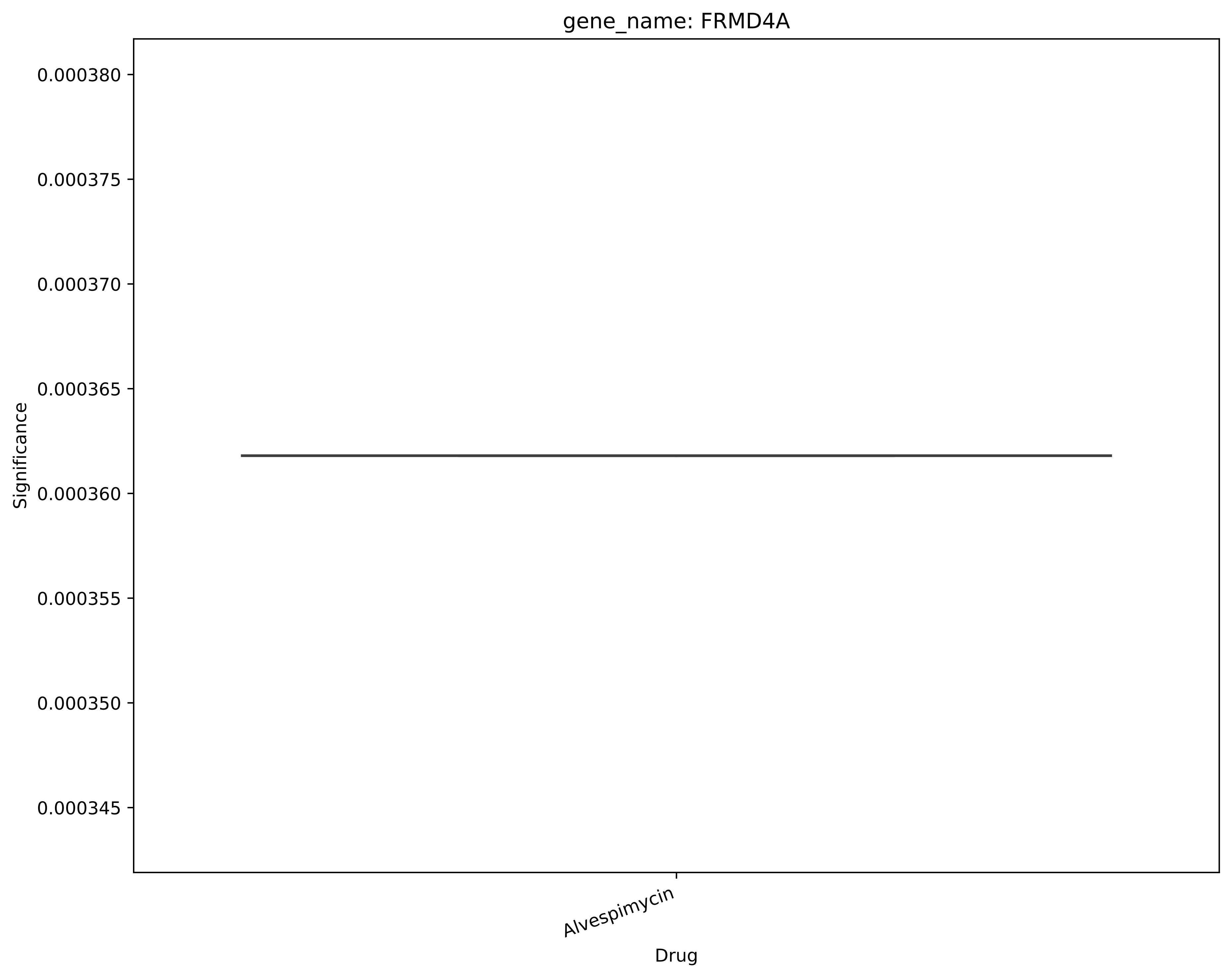

gene_name: FRMD4A
Significance
0.000380
0.000375
0.000370
0.000365
0.000360
0.000355
0.000350
0.000345
Alvespimycin
Drug

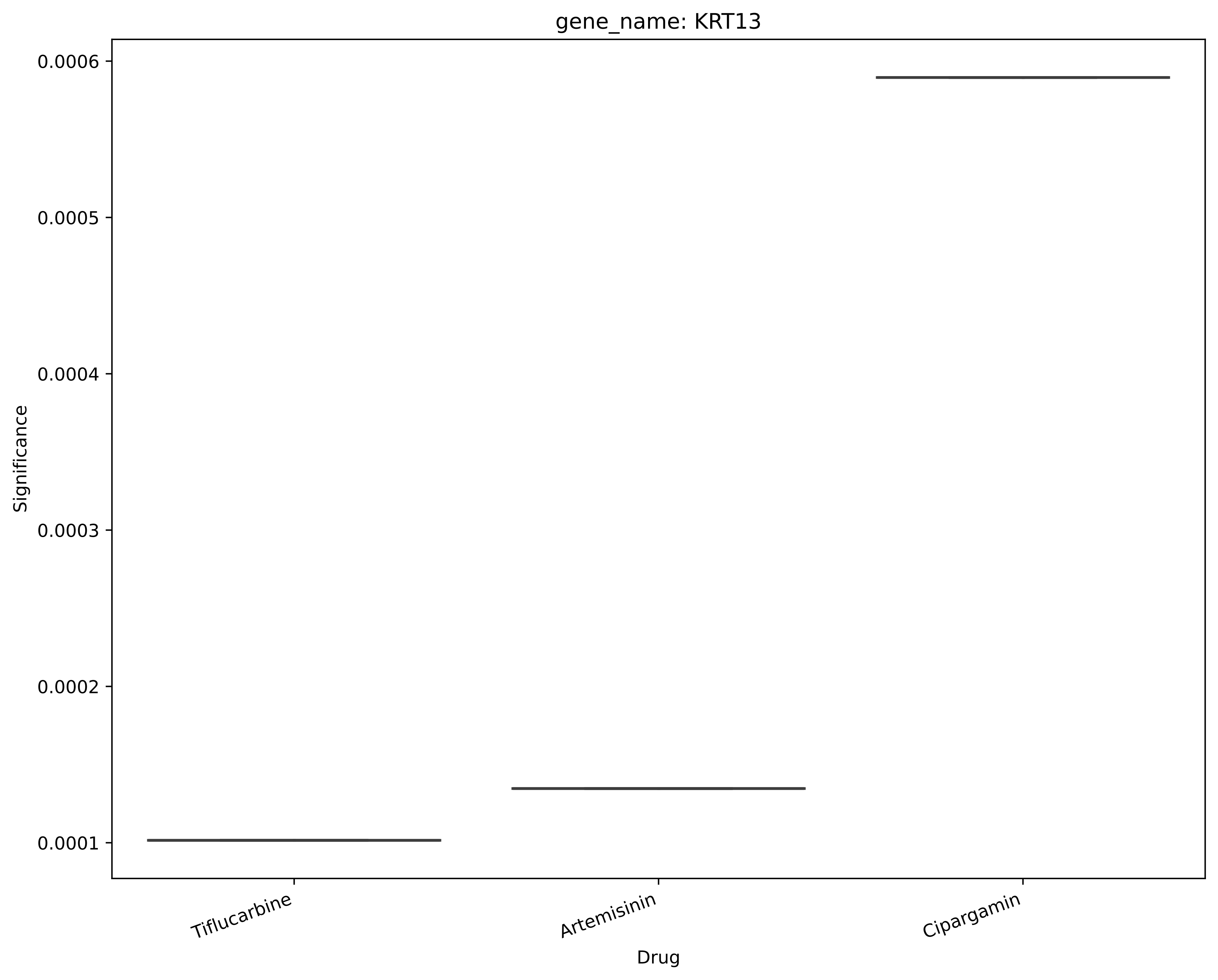
gene_name: KRT13
Significance
0.0006
0.0005
0.0004
0.0003
0.0002
0.0001
Tiflucarbine
Artemisinin
Cipargamin
Drug

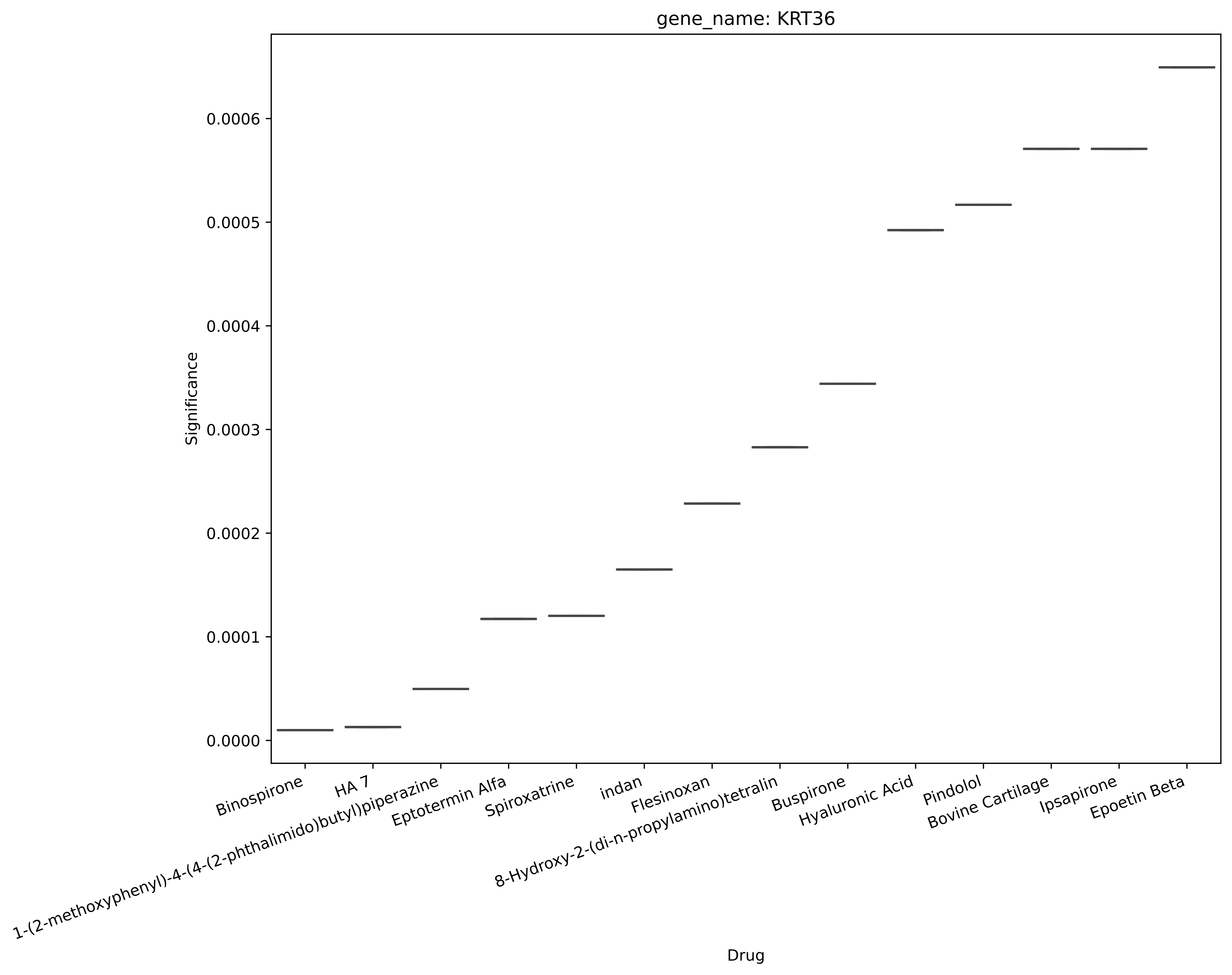
gene_name: KRT36
Significance
0.0000
0.0001
0.0002
0.0003
0.0004
0.0005
0.0006
Binospirone
HA 7
1-(2-methoxyphenyl)-4-(4-(2-phthalimido)butyl)piperazine
Eptotermin Alfa
Spiroxatrine
indan
Flesinoxan
8-Hydroxy-2-(di-n-propylamino)tetralin
Buspirone
Hyaluronic Acid
Pindolol
Bovine Cartilage
Ipsapirone
Epoetin Beta
Drug

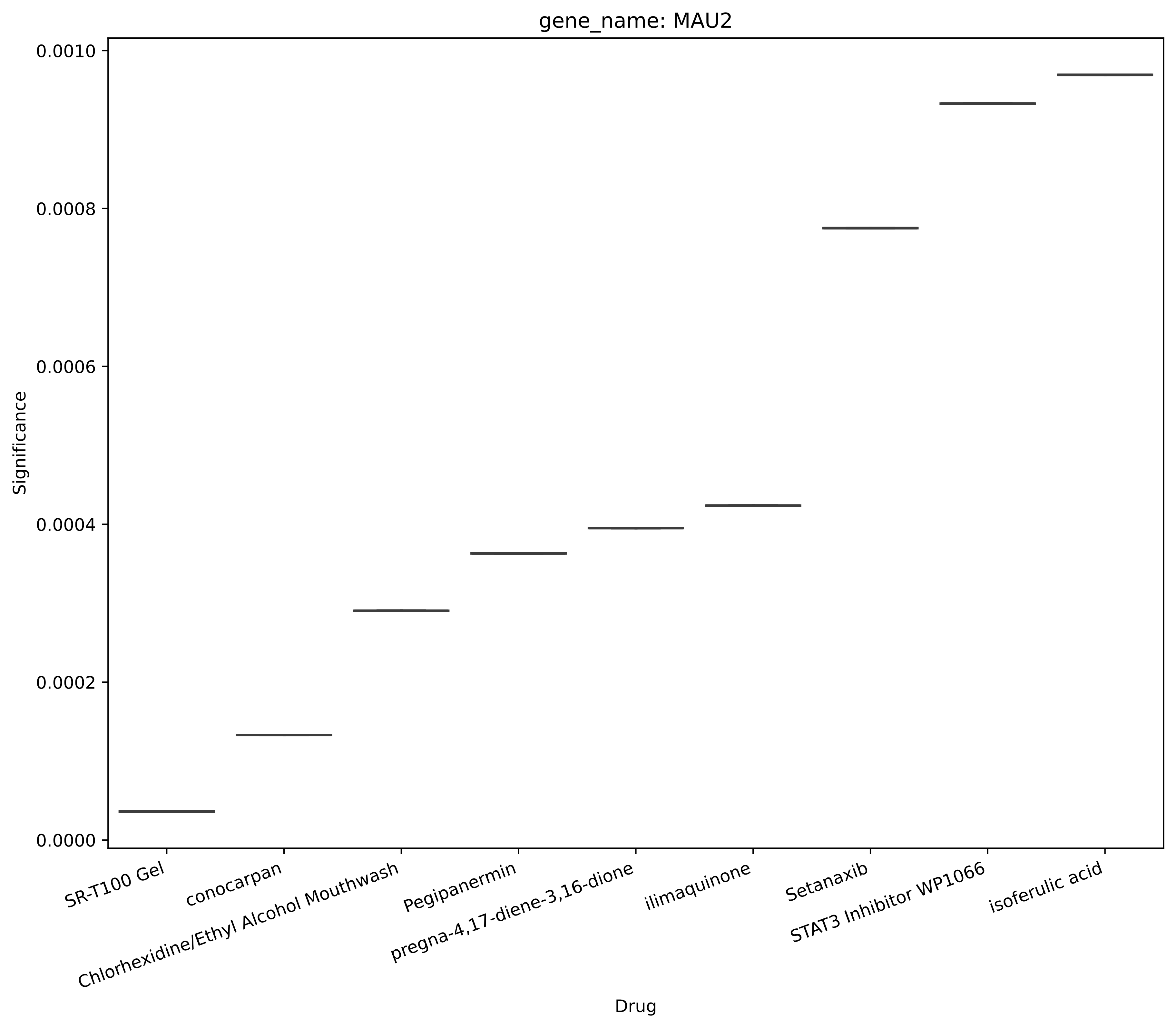

gene_name: MAU2
Significance
0.0000
0.0002
0.0004
0.0006
0.0008
0.0010
Drug
SR-T100 Gel
conocarpan
Chlorhexidine/Ethyl Alcohol Mouthwash
Pegipanermin
pregna-4,17-diene-3,16-dione
ilimaquinone
Setanaxib
STAT3 Inhibitor WP1066
isoferulic acid

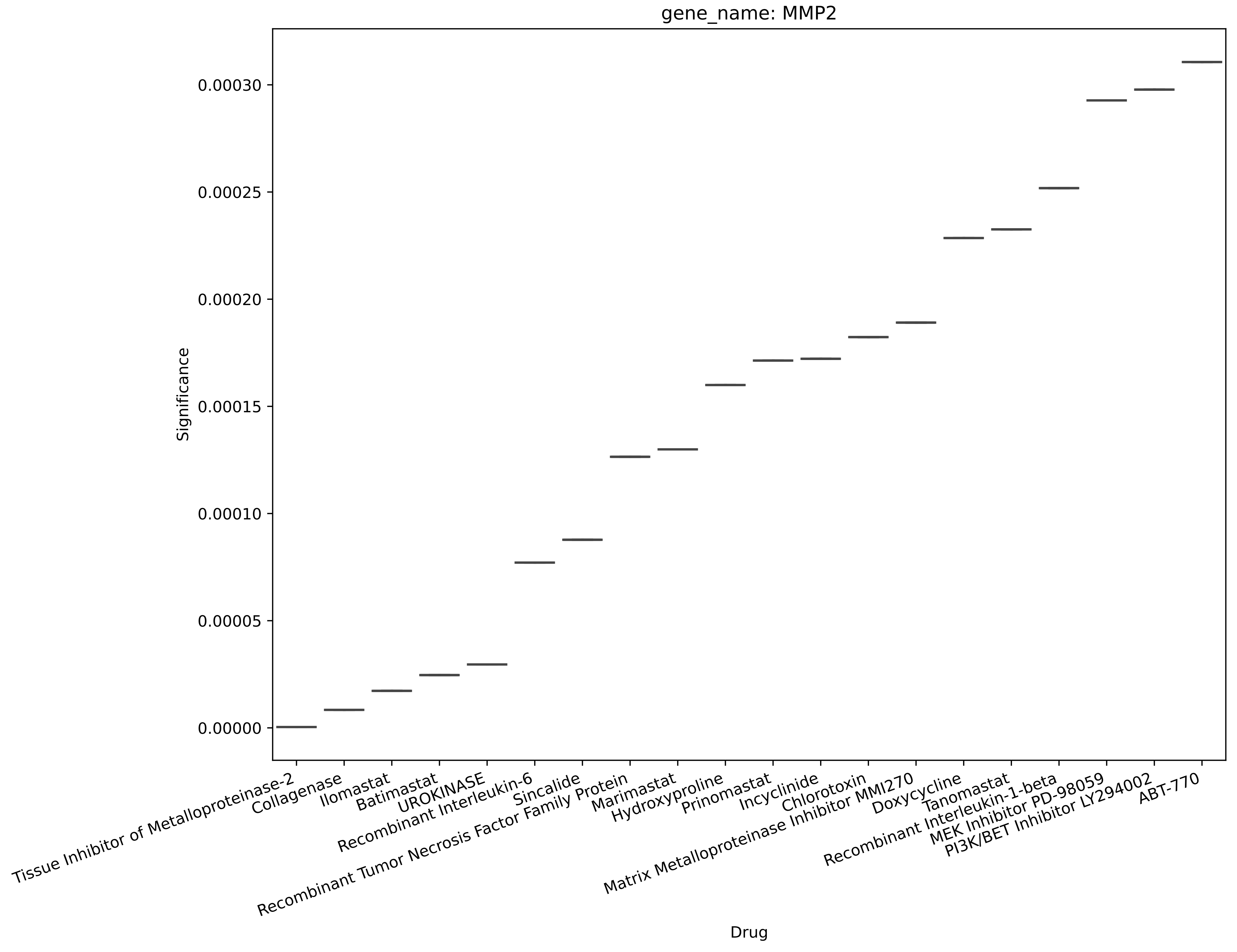
gene_name: MMP2
Significance
0.00030
0.00025
0.00020
0.00015
0.00010
0.00005
0.00000
Tissue Inhibitor of Metalloproteinase-2
Collagenase
Ilomastat
Batimastat
UROKINASE
Recombinant Interleukin-6
Sincalide
Recombinant Tumor Necrosis Factor Family Protein
Marimastat
Hydroxyproline
Prinomastat
Incyclinide
Chlorotoxin
Matrix Metalloproteinase Inhibitor MMI270
Doxycycline
Tanomastat
Recombinant Interleukin-1-beta
MEK Inhibitor PD-98059
PI3K/BET Inhibitor LY294002
ABT-770
Drug

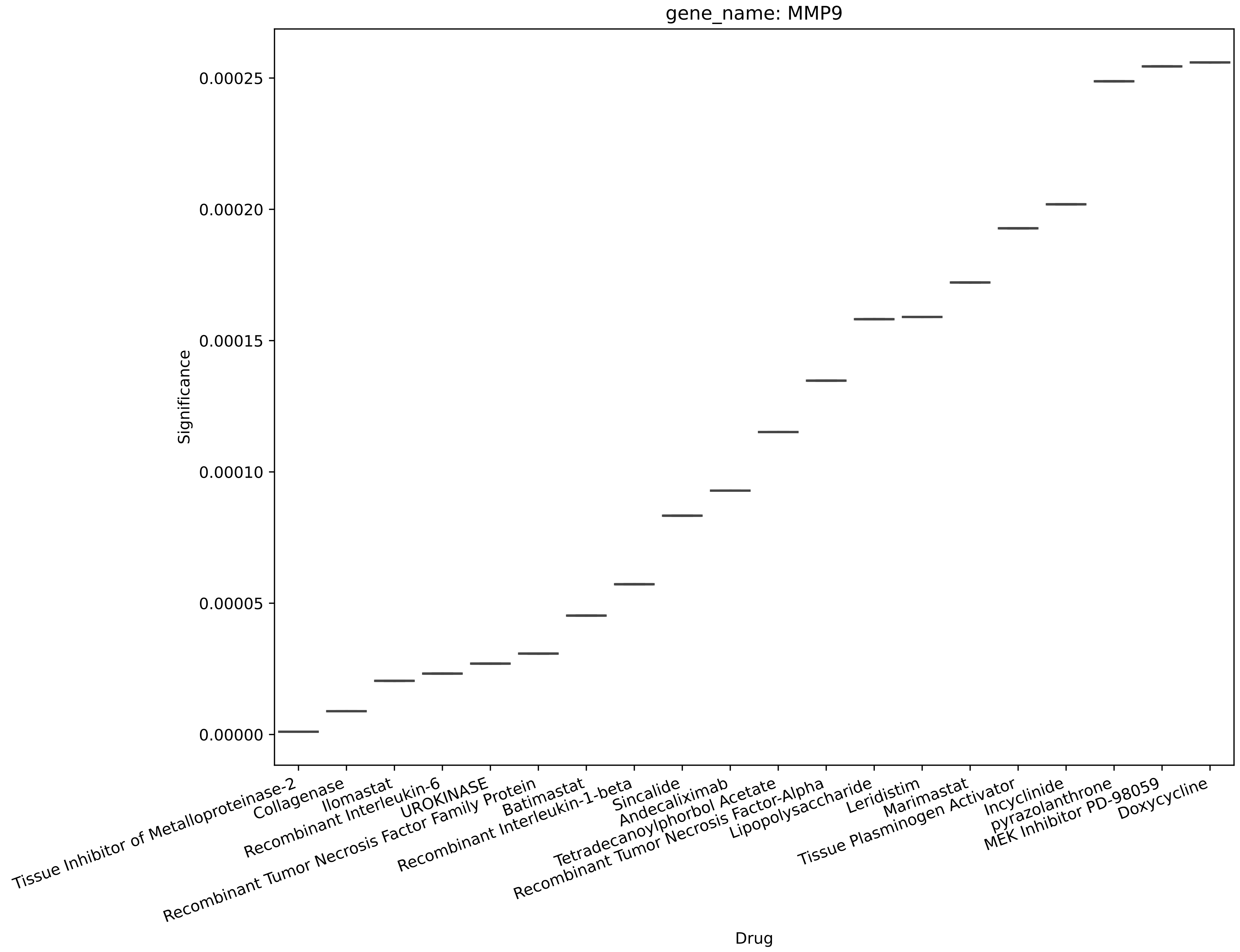
gene_name: MMP9
Significance
0.00025
0.00020
0.00015
0.00010
0.00005
0.00000
Tissue Inhibitor of Metalloproteinase-2
Collagenase
Ilomastat
Recombinant Interleukin-6
UROKINASE
Recombinant Tumor Necrosis Factor Family Protein
Batimastat
Recombinant Interleukin-1-beta
Sincalide
Andecaliximab
Tetradecanoylphorbol Acetate
Recombinant Tumor Necrosis Factor-Alpha
Lipopolysaccharide
Leridistim
Marimastat
Tissue Plasminogen Activator
Incyclinide
pyrazolanthrone
MEK Inhibitor PD-98059
Doxycycline
Drug

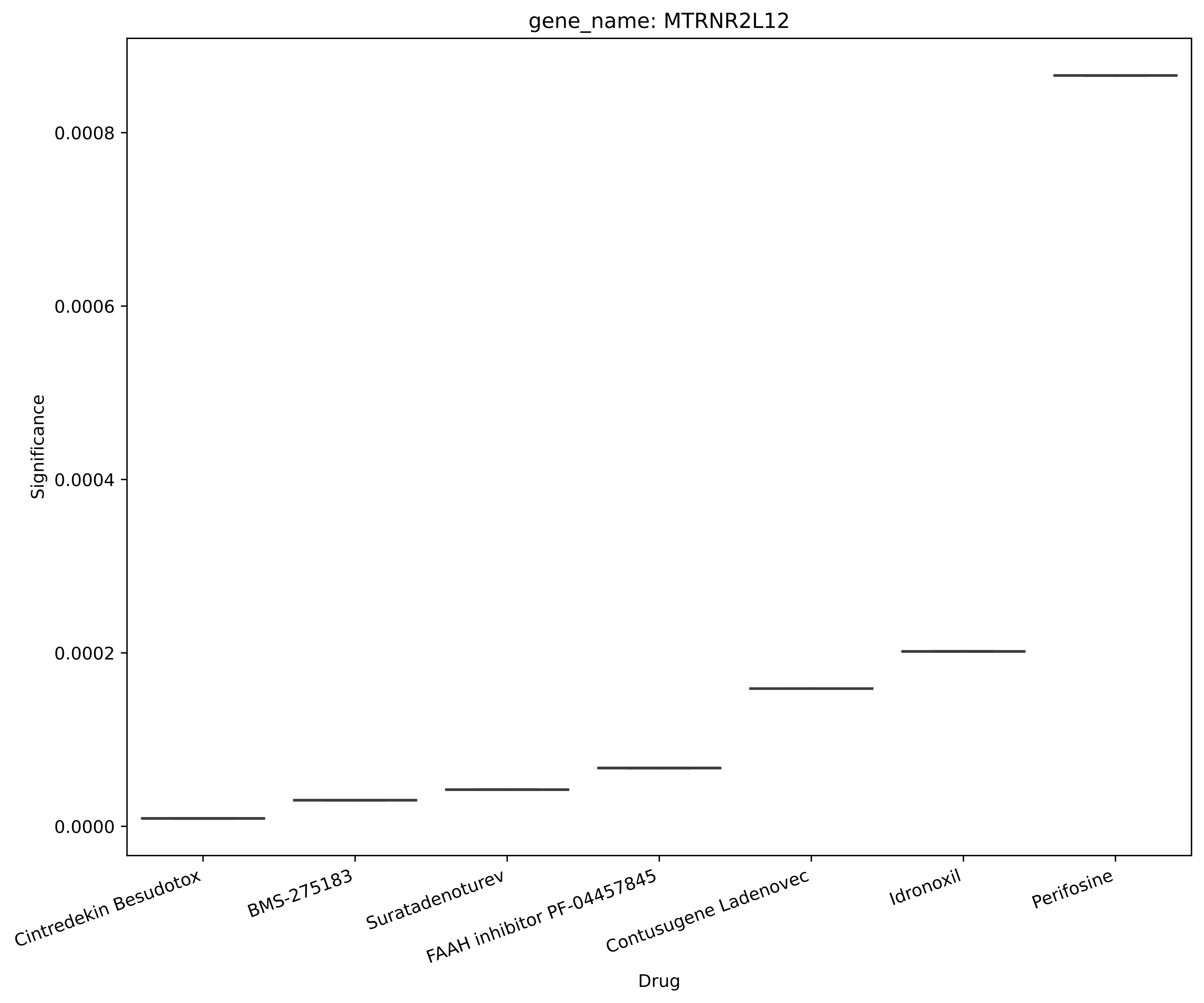

gene_name: MTRNR2L12
Significance
0.0000
0.0002
0.0004
0.0006
0.0008
Cintredekin Besudotox
BMS-275183
Suratadenoturev
FAAH inhibitor PF-04457845
Contusugene Ladenovec
Idronoxil
Perifosine
Drug

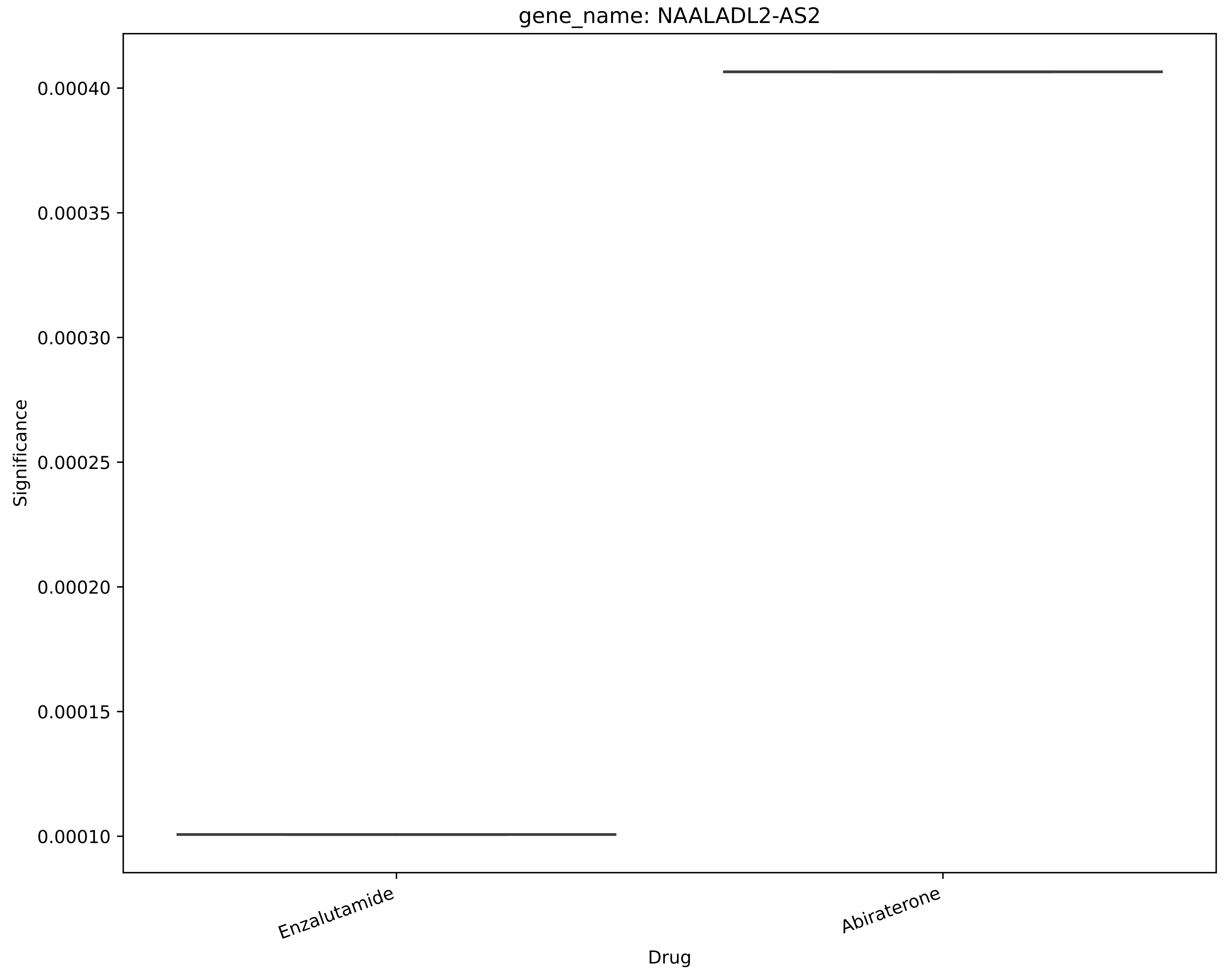

gene_name: NAALADL2-AS2
Significance
0.00040
0.00035
0.00030
0.00025
0.00020
0.00015
0.00010
Enzalutamide
Abiraterone
Drug

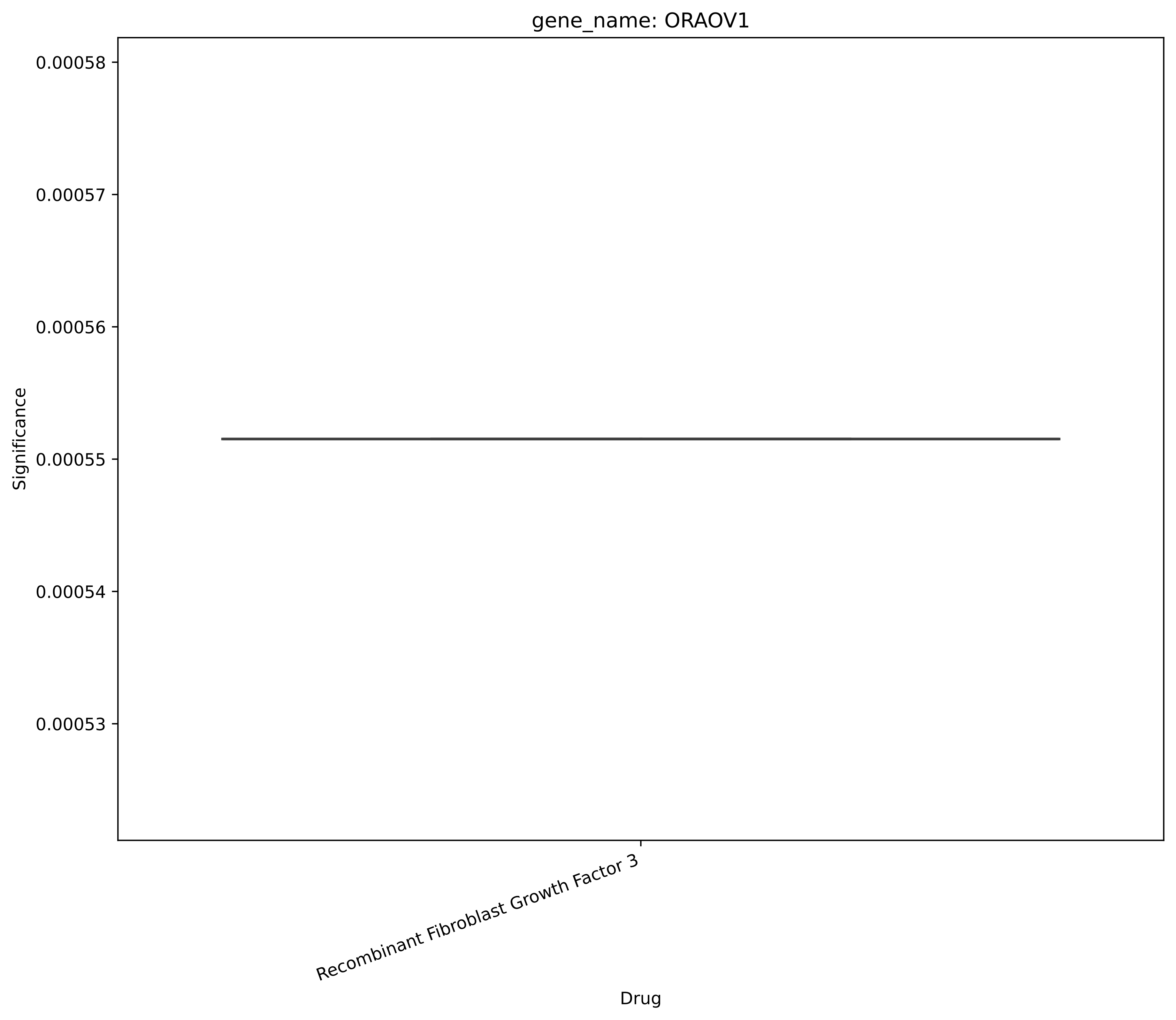

gene_name: ORAOV1
0.00058
0.00057
0.00056
0.00055
0.00054
0.00053
Significance
Recombinant Fibroblast Growth Factor 3
Drug

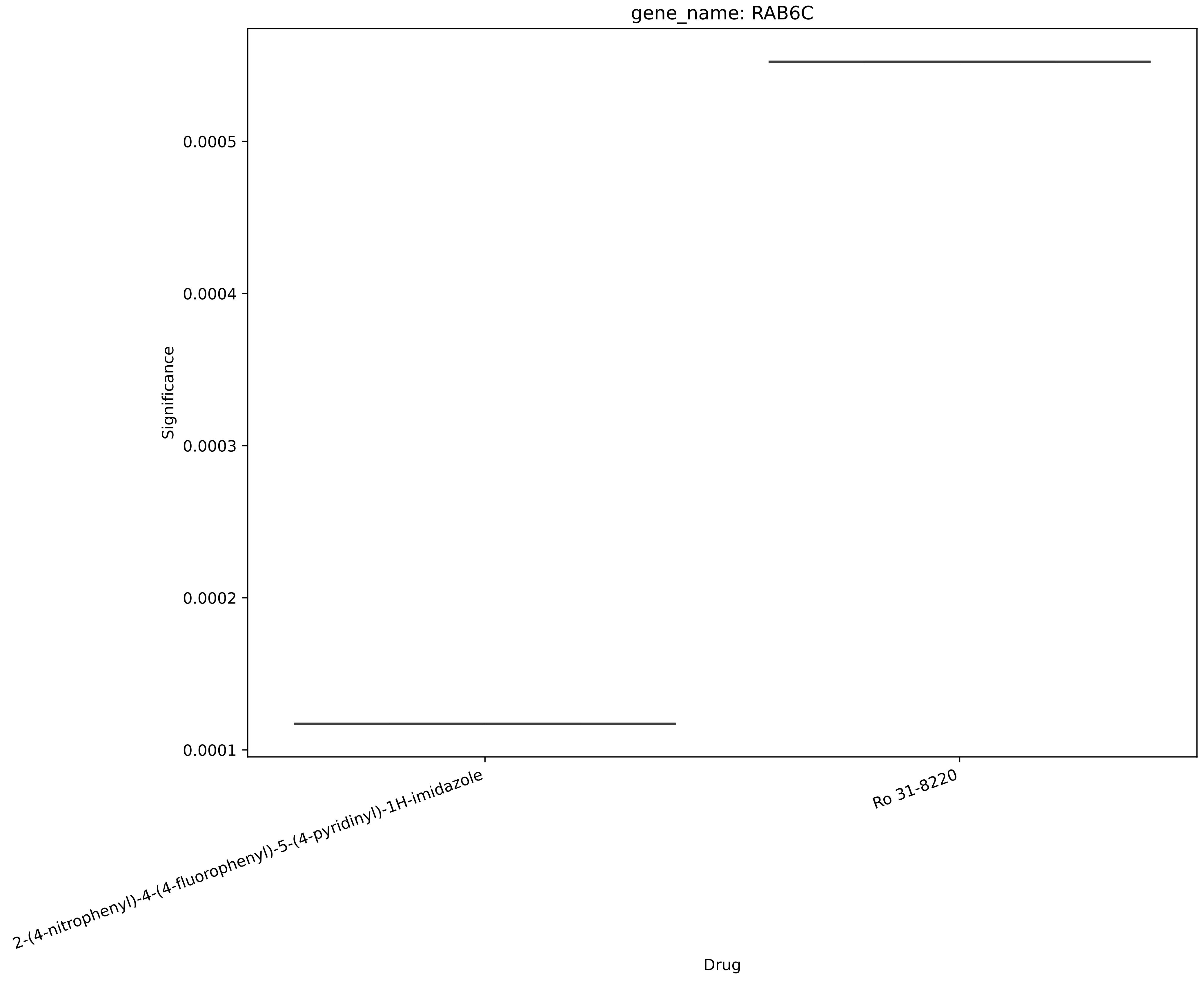
gene_name: RAB6C
Significance
0.0005
0.0004
0.0003
0.0002
0.0001
2-(4-nitrophenyl)-4-(4-fluorophenyl)-5-(4-pyridinyl)-1H-imidazole
Ro 31-8220
Drug

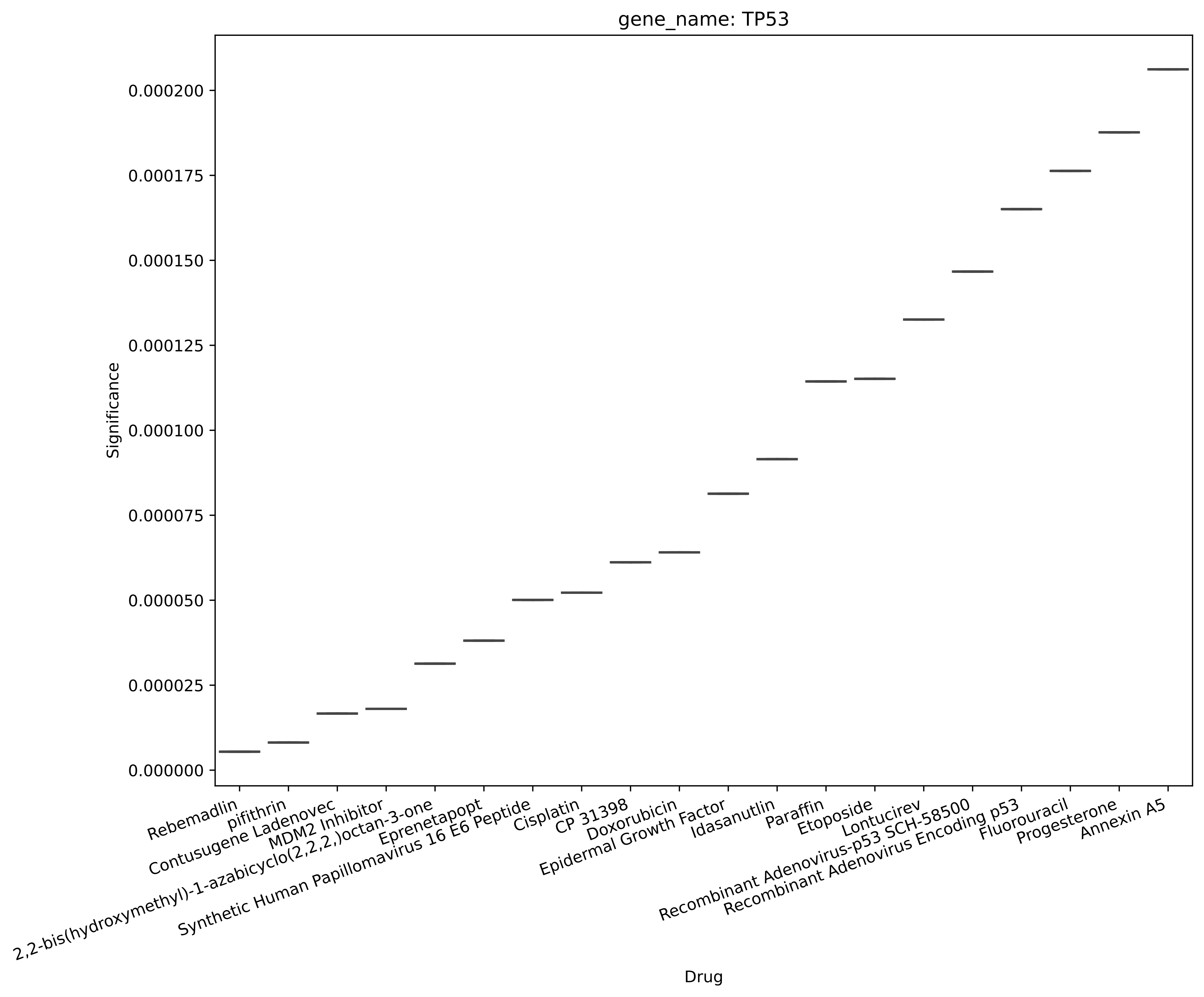

gene_name: TP53
Significance
0.000000
0.000025
0.000050
0.000075
0.000100
0.000125
0.000150
0.000175
0.000200
Rebemadlin
pifithrin
Contusugene Ladenovec
MDM2 Inhibitor
2,2-bis(hydroxymethyl)-1-azabicyclo(2,2,2,)octan-3-one
Eprenetapopt
Synthetic Human Papillomavirus 16 E6 Peptide
Cisplatin
CP 31398
Doxorubicin
Epidermal Growth Factor
Idasanutlin
Paraffin
Etoposide
Lontucirev
Recombinant Adenovirus-p53 SCH-58500
Recombinant Adenovirus Encoding p53
Fluorouracil
Progesterone
Annexin A5
Drug

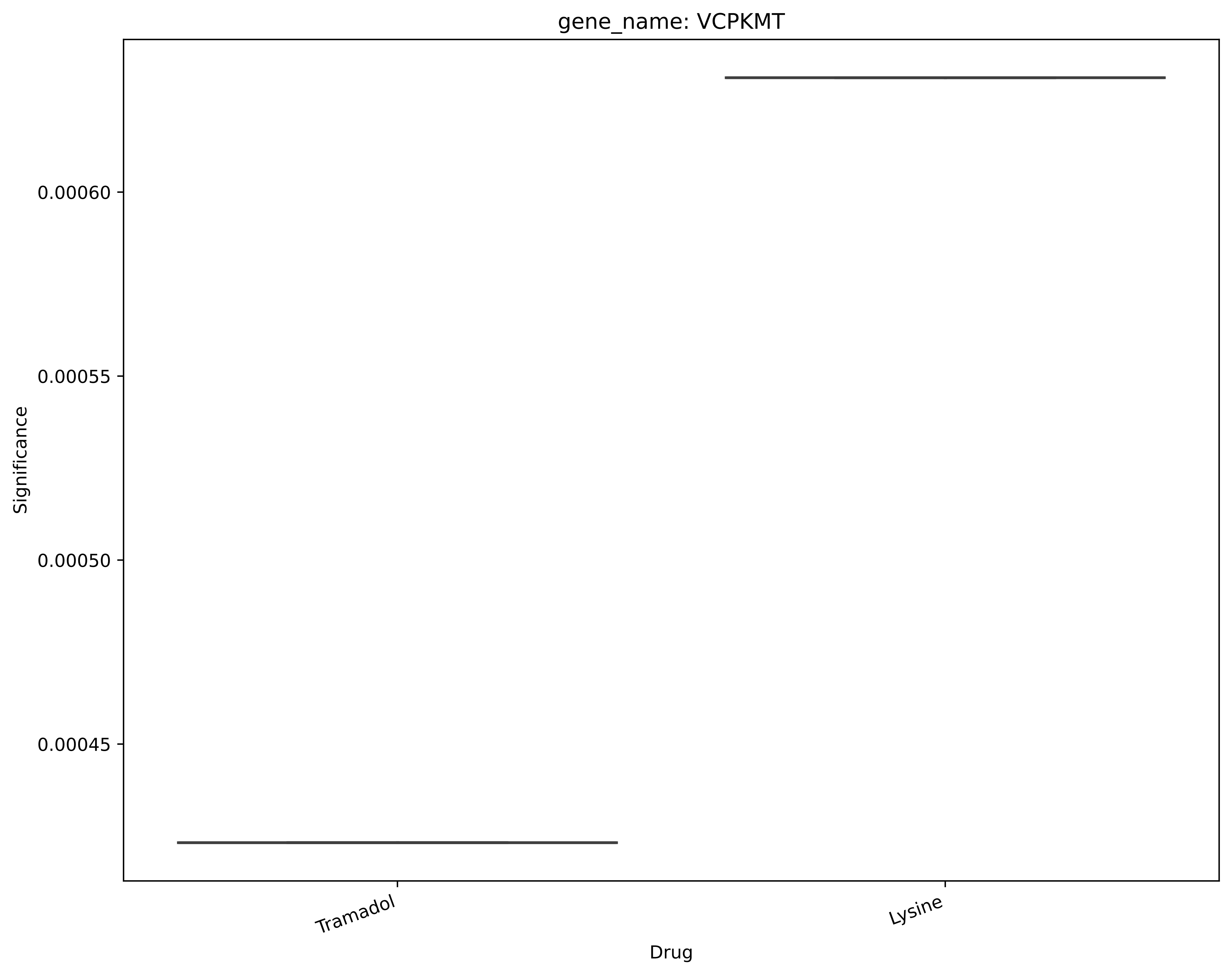
gene_name: VCPKMT
0.00060
0.00055
0.00050
0.00045
Significance
Tramadol
Lysine
Drug

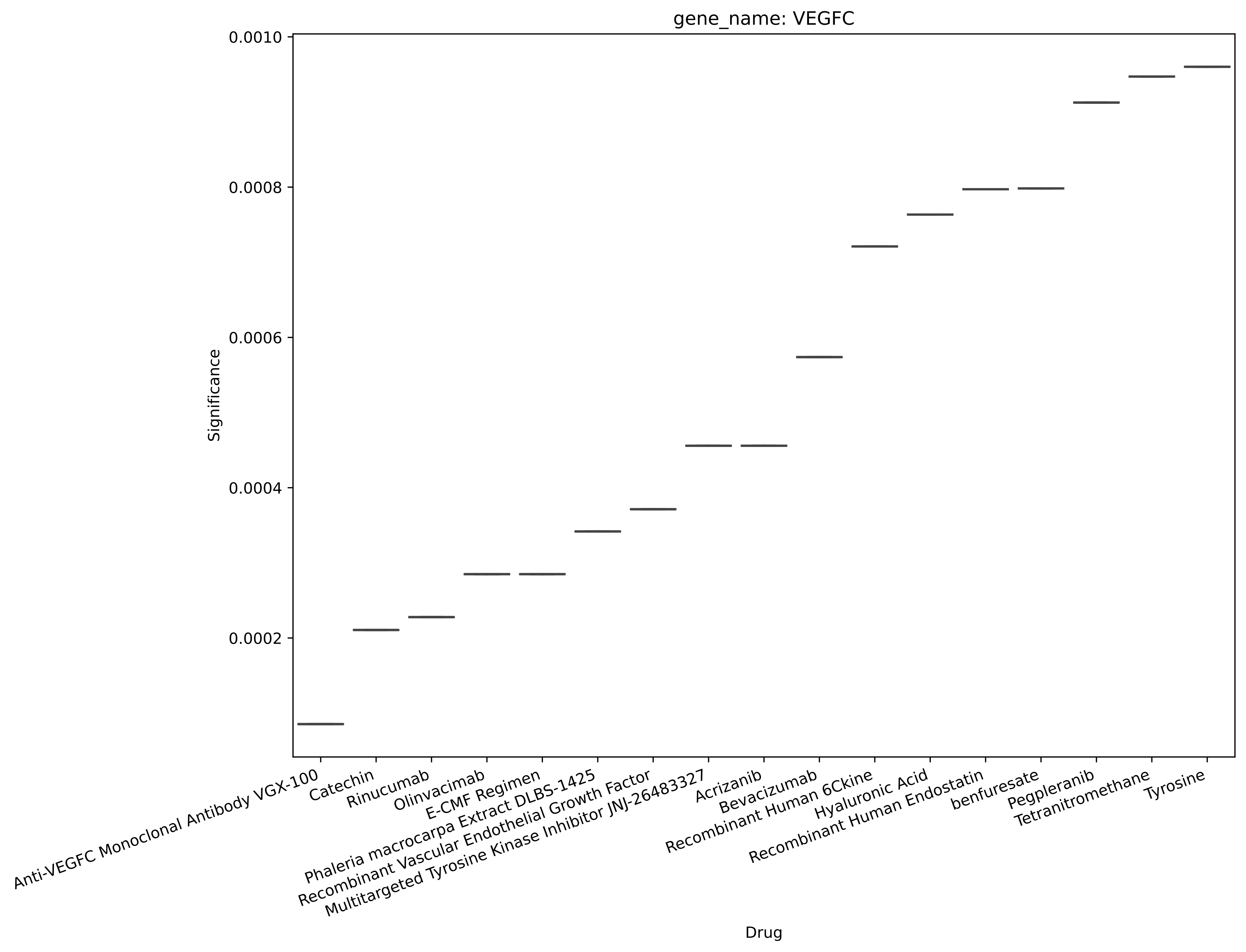

gene_name: VEGFC
Significance
0.0010
0.0008
0.0006
0.0004
0.0002
Drug
Anti-VEGFC Monoclonal Antibody VGX-100
Catechin
Rinucumab
Olinvacimab
E-CMF Regimen
Phaleria macrocarpa Extract DLBS-1425
Recombinant Vascular Endothelial Growth Factor
Multitargeted Tyrosine Kinase Inhibitor JNJ-26483327
Acrizanib
Bevacizumab
Recombinant Human 6Ckine
Hyaluronic Acid
Recombinant Human Endostatin
benfuresate
Pegpleranib
Tetranitromethane
Tyrosine

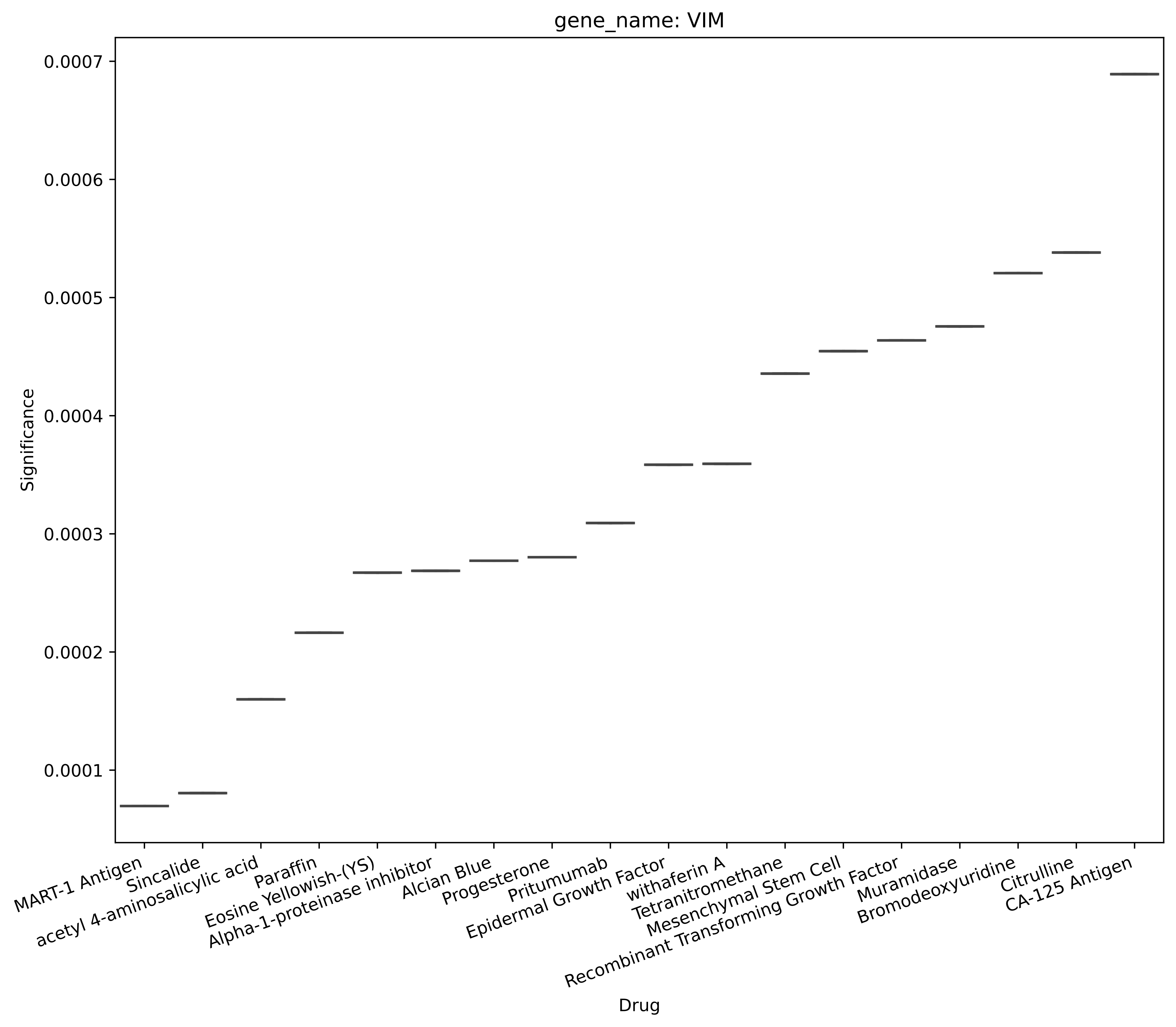

gene_name: VIM
Significance
0.0007
0.0006
0.0005
0.0004
0.0003
0.0002
0.0001
Drug
MART-1 Antigen
Sincalide
acetyl 4-aminosalicylic acid
Paraffin
Eosine Yellowish-(YS)
Alpha-1-proteinase inhibitor
Alcian Blue
Progesterone
Pritumumab
Epidermal Growth Factor
withaferin A
Tetranitromethane
Mesenchymal Stem Cell
Recombinant Transforming Growth Factor
Muramidase
Bromodeoxyuridine
Citrulline
CA-125 Antigen

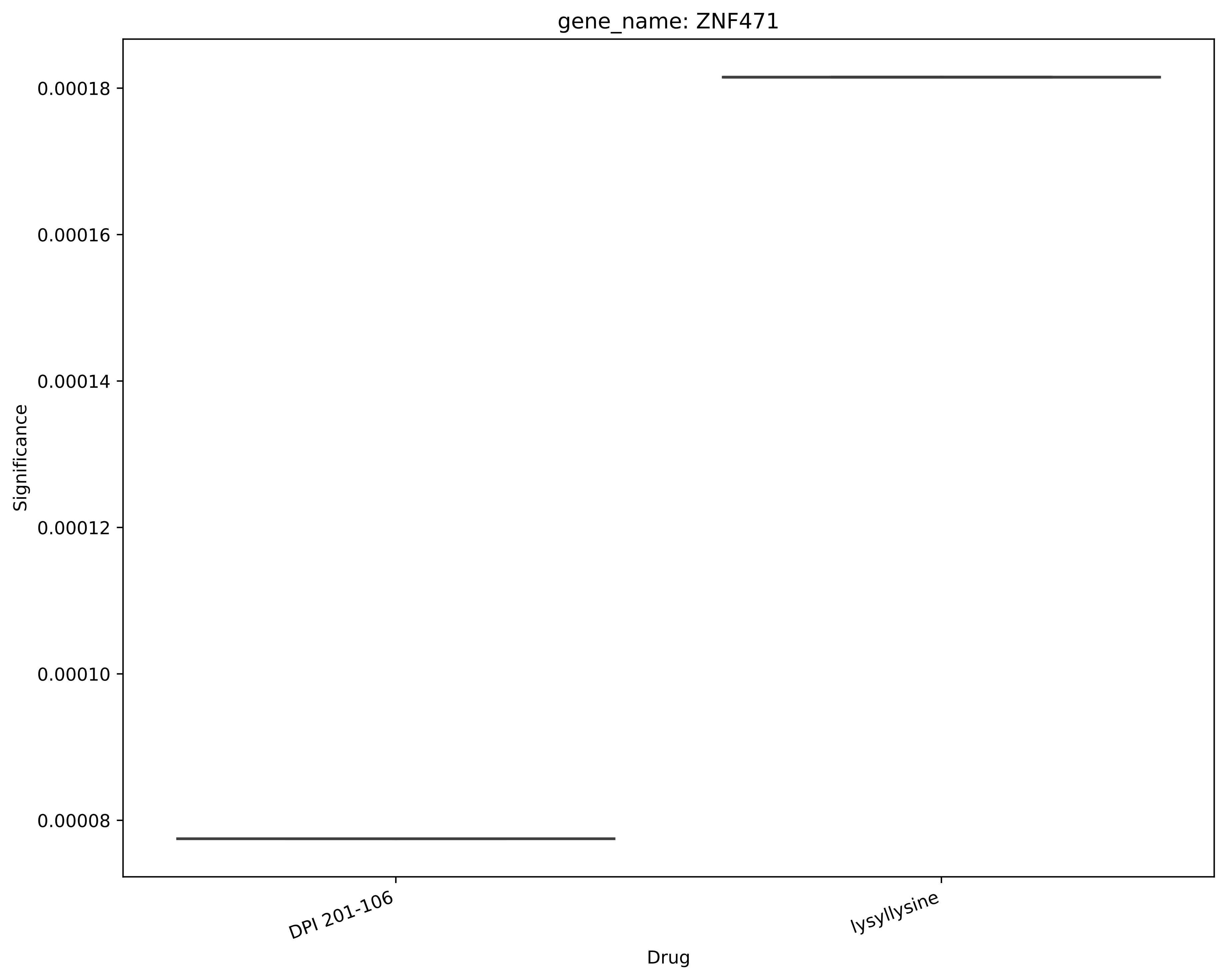
gene_name: ZNF471
Significance
0.00018
0.00016
0.00014
0.00012
0.00010
0.00008
DPI 201-106
lysyllysine
Drug
